# A brain-body feedback loop driving HPA-axis dysfunction in breast cancer

**DOI:** 10.1101/2024.09.13.612923

**Authors:** Adrian Gomez, Yue Wu, Chao Zhang, Leah Boyd, Tse-Luen Wee, Joseph Gewolb, Corina Amor, Lucas Cheadle, Jeremy C. Borniger

## Abstract

Breast cancer patients often exhibit disrupted circadian rhythms in circulating glucocorticoids (GCs), such as cortisol. This disruption correlates with reduced quality of life and higher cancer mortality^1–3^. The exact cause of this phenomenon — whether due to treatments, stress, age, co-morbidities, lifestyle factors, or the cancer itself remains unclear. Here, we demonstrate that primary breast cancer alone blunts host GC rhythms by disinhibiting neurons in the hypothalamus, and that circadian phase-specific neuromodulation of these neurons can attenuate tumor growth by enhancing anti-tumor immunity. We find that mice with mammary tumors exhibit blunted GC rhythms before tumors are palpable, alongside increased activity in paraventricular hypothalamic neurons expressing corticotropin-releasing hormone (i.e., PVN^CRH^ neurons). Tumor-bearing mice have fewer inhibitory synapses contacting PVN^CRH^ neurons and reduced miniature inhibitory post-synaptic current (mIPSC) frequency, leading to net excitation. Tumor-bearing mice experience impaired negative feedback on GC production, but adrenal and pituitary gland functions are largely unaffected, indicating that alterations in PVN^CRH^ neuronal activity are likely a primary cause of hypothalamic-pituitary-adrenal (HPA) axis dysfunction in breast cancer. Using chemogenetics (hM3Dq) to stimulate PVN^CRH^ neurons at different circadian phases, we show that stimulation just before the light-to-dark transition restores normal GC rhythms and reduces tumor progression. These mice have significantly more effector T cells (CD8+) within the tumor than non-stimulated controls, and the anti-tumor effect of PVN^CRH^ neuronal stimulation is absent in mice lacking CD8+ T cells. Our findings demonstrate that breast cancer distally regulates neurons in the hypothalamus that control output of the HPA axis and provide evidence that therapeutic targeting of these neurons could mitigate tumor progression.

## Main

Breast cancer remains one of the most ubiquitous and life-threatening conditions among women worldwide^4^. Although we have made substantial progress in early detection and therapy, individuals with breast cancer still exhibit a diverse array of systemic problems that can significantly influence their prognoses (e.g., fatigue and sleep difficulties, cognitive impairments)^3,5–7^. A lesser-known predictor of breast cancer mortality and quality of life is the blunting of circadian rhythms in circulating glucocorticoids (GCs) during cancer progression. Specifically, patients with ‘flat’ GC slopes (indicating little difference between morning and evening GC concentrations) experience the highest levels of fatigue, exhibit reduced numbers and suppressed activity of circulating natural killer (NK) cells, and impaired T cell-mediated immunity, while also succumbing significantly earlier to cancer than patients that maintain robust circadian GC rhythms^1–3,8–10^. This phenomenon is also evident in lung^11^, colorectal^12^, and ovarian^13^ cancer, indicating that there may be shared mechanisms driving GC rhythm dysfunction across cancers.

Maintenance of robust GC rhythms influences immune cell trafficking^14^, systemic energy balance^15^, and entrainment of cellular circadian clocks^16^, among other functions^17^. This has given rise to the idea that the oscillatory pattern of GC expression, with the peak at the onset of the active phase and the nadir in the late inactive phase, is essential for daily physiological needs^17^. Glucocorticoid rhythms are controlled primarily via concerted actions among the hypothalamus (specifically the suprachiasmatic nucleus (SCN) and the paraventricular nucleus (PVN)), anterior pituitary, and adrenal cortex (i.e., the HPA axis)^18,19^. Despite ample clinical evidence, we still do not understand how GC rhythm dysfunction is linked to breast cancer outcomes.

### Breast cancer ‘blunts’ circadian rhythms in endogenous glucocorticoids in mice

A major obstacle to understanding the relationship between host glucocorticoid rhythms and cancer outcomes is the numerous co-variates that can influence GC concentrations (e.g., cancer treatment regimen, psychological stress, or age). This has made it difficult to determine whether cancer alone, independent of other factors, can influence host GC rhythms. To resolve this, we turned to mouse models that allow us to control most of these factors and give us access to GC concentrations throughout the circadian cycle. We collected fecal samples every six hours across the day (Zeitgeber Time (ZT) 0, 6, 12, 18, 24) at baseline and then weekly throughout the course of tumor progression in the orthotopic syngeneic EO771 breast cancer model^20^ (**Fig. 1d**). Measuring GCs in fecal samples allowed us to sample GCs without causing additional stress to the animals, which may confound the results^21^. We observed a ∼33-55% reduction in glucocorticoid concentrations during cancer progression, starting just one week following tumor cell injection and continuing throughout the experiment (4 weeks) (**Fig. 1e, f and Extended Data Fig. 1b**). This finding was absent in sham-injected mice as robust GC rhythms were maintained during consecutive weeks of GC measurement (**Fig. 1a-c and Extended Data Fig. 1a**). Interestingly, tumor size was not correlated with glucocorticoid concentrations (**Extended Data Fig. 1e, f**), suggesting the presence of tumor cells, independent of tumor weight, is sufficient to dampen glucocorticoid rhythms. Furthermore, the ZT12 timepoint, where GC rhythms peak (i.e., acrophase), was most affected by the presence of the tumor (**Fig. 1e**). This effect is reflected in the blunted slope of the rhythm (**Fig. 1e, inset**), a similar phenomenon to that observed in human patients^1,2^. We repeated this experiment in an autochthonous breast cancer model (c57bl6 MMTV-PyMT) where mammary tumors develop spontaneously after approximately 92 days of age^22^ (**Fig. 1g**). We observed that blunting of the rhythm occurs only after spontaneous development of palpable breast tumors, with specific reductions in fecal GCs at ZT6 and ZT12 (**Fig. 1h-j**). To determine whether blunted rhythms persist following removal of the tumor, we surgically resected the primary tumor and continued to monitor glucocorticoid rhythms over the next two months (**Fig. 1k, l**). Glucocorticoid rhythms remained blunted for at least 8 weeks following removal of the tumor, with no observable signs of metastasis (**Fig. 1m, n and Extended Data Fig. 1g, h**), suggesting that tumor-induced changes in HPA axis function can endure well past tumor removal and recovery from surgical resection.

**Figure 1.**
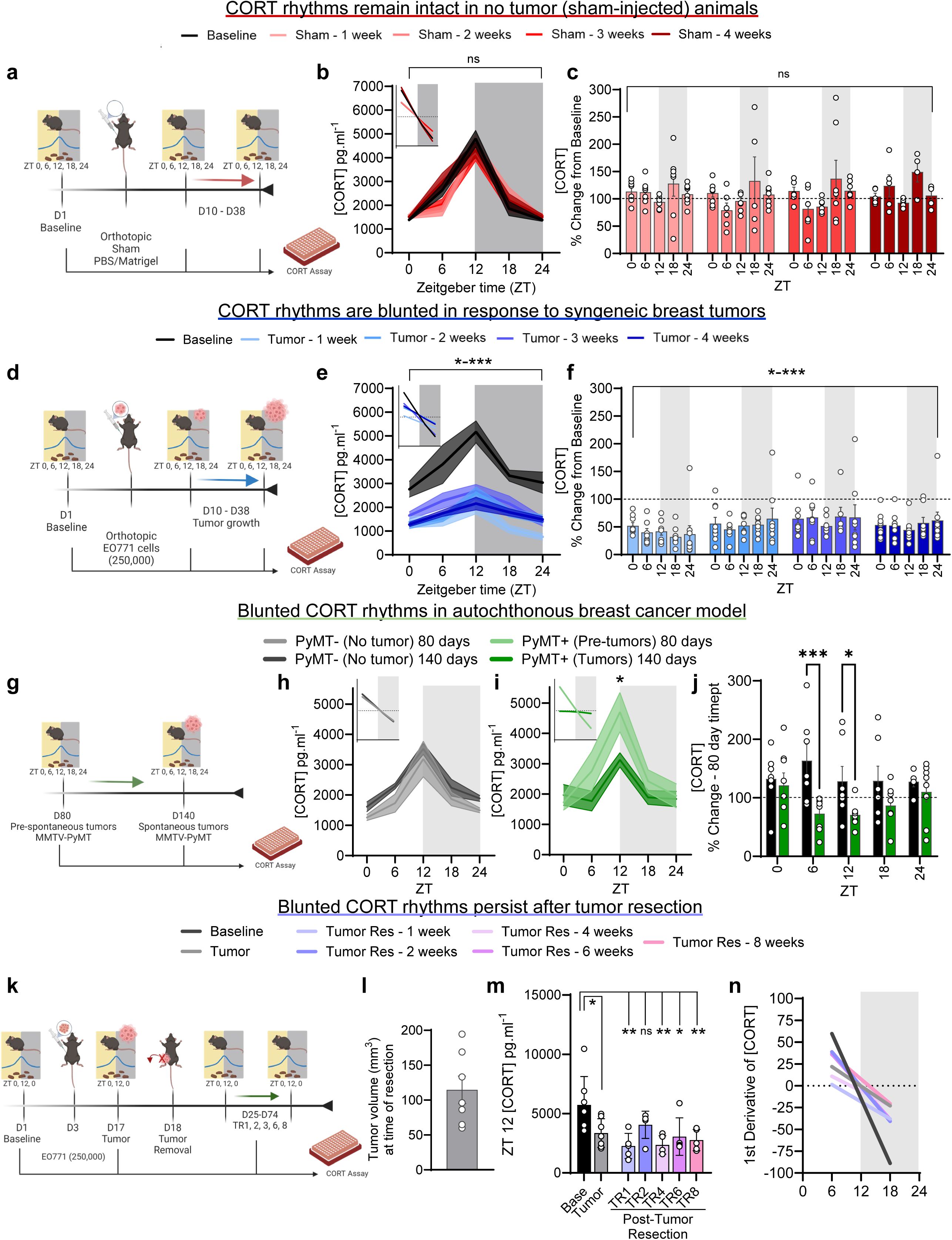
Breast cancer blunts host glucocorticoid circadian rhythms. **a,** Schematic depicting orthotopic injections of PBS/Matrigel (sham) in female WT mice and subsequent fecal boli collections for corticosterone measurements during progressive weeks. **b,** Fecal corticosterone levels across the day (6 hr epochs) in sham-injected mice during baseline and progressive weeks. Inset represents first derivative of corticosterone measurements, corresponding to panel **b**, depicting slope of the tangent lines in sham-injected mice during baseline and progressive weeks. n = 5-8 mice / timepoint. **c,** Fecal corticosterone levels in sham-injected expressed as a percent change from baseline. n = 5-8 mice / timepoint. **d,** Schematic depicting orthotopic injections of EO771 cells (250,000) in female WT mice and subsequent fecal boli collections for corticosterone measurements during tumor progression. **e,** Fecal corticosterone levels across the day (6 hr epochs) in EO771-injected mice during baseline and tumor progression. Inset represents first derivative of corticosterone measurements, corresponding to panel **e**, depicting slope of the tangent lines in EO771-injected mice during baseline and tumor progression. n=9-11 mice / timepoint. **f,** Fecal corticosterone levels in EO771-injected expressed as a percent change from baseline. n = 9-11 mice / timepoint. **g,** Schematic illustrating fecal corticosterone collection from the MMTV-PyMT mouse strain which spontaneous develops palpable mammary tumors starting around 90 days. **h,** Fecal corticosterone levels across the day (6 hr epochs) in MMTV-PyMT-mice at 80 and 140 days of age who have not developed palpable tumors. Inset depicts the first derivative of corticosterone measurements from panel (**h**). n = 5-7 mice / timepoint for 80 days; n=9 / timepoint for 140 days. **i,** Fecal corticosterone levels across the day (6 hr epochs) in MMTV-PyMT+ mice at 80 and 140 days of age who have developed palpable tumors (day 140). Inset depicts the first derivative of corticosterone measurements from panel (**i**). n=8 mice / timepoint for 80 and 140 days. **j,** Fecal corticosterone levels in MMTV-PyMT mice expressed as a percent change from baseline (day 80). **k,** Schematic depicting tumor resection and subsequent corticosterone measurements. **l,** Tumor volume at time of tumor resection. n = 7 mice. **m,** Corticosterone measurements at ZT12 in mice at baseline, tumor, and “tumor-free” timepoints. n = 7 mice for baseline and tumor timepoints; n=5 for post-tumor resection weeks. Note: A tumor was detected in 2 of the mice 1 week after resection, and were removed from subsequent “tumor-free” measurements. **n,** First derivative of corticosterone measurements in mice at baseline, tumor, and “tumor-free” timepoints. All data represented as mean ± SEM. \**p* < 0.05, \*\**p* < 0.01, \*\*\**p* < 0.001; two-way repeated-measures ANOVA followed by separate one-way ANOVAs with Bonferroni multiple comparisons (**b, c, e, f, h-j, m**).

We next assessed how each component of the HPA axis functionally changes during breast cancer progression (**Extended Data Fig. 2a, b**). To examine adrenal responsiveness to hypothalamic-pituitary output, we challenged mice with exogenous adrenocorticotropic hormone (ACTH; 5 mg kg^-1^ or 50 mg kg^-1^), which normally elicits glucocorticoid release via ACTH receptor (MC2R) activation in the adrenal cortex. Tumor-bearing mice had decreased basal corticosterone concentrations (as in **Fig. 1e**), but produced similar amounts of corticosterone in response to ACTH, suggesting that adrenal sensitivity to ACTH is intact (**Extended Data Fig. 2e**). Consistent with this, we found no changes in the expression of *Mc2r* or steroidogenic enzymes (e.g., *Cyp11b1, Hsd3b1,* and *StAR*) in the adrenal glands of tumor-bearing mice (**Extended Data Fig. 2d**), despite an increase in adrenal mass (**Extended Data Fig. 2c**). Within the pituitary, we saw no differences in expression of *Pomc* (the precursor of ACTH) or the glucocorticoid receptor (*Nr3c1*) in tumor-bearing mice (**Extended Data Fig. 2h**), suggesting that breast cancer disrupts endogenous glucocorticoid rhythms, but that steroidogenic functions of the pituitary and adrenal glands remain largely unaffected.

Glucocorticoid production is also regulated via negative feedback loops involving glucocorticoid (GRs) and mineralocorticoid (MRs) receptors within the pituitary and centrally in the PVN^23^. To assess negative feedback in mice with breast tumors, we injected animals with the synthetic glucocorticoid dexamethasone (DEX, 20 mg kg^-1^, IP), a potent GR agonist that decreases endogenous glucocorticoid concentrations via negative feedback suppression^24^ (**Extended Data Fig. 2b**). As expected, we observed a decrease in glucocorticoids in response to DEX in non-tumor-bearing mice; however, in mice harboring EO771 tumors, DEX failed to decrease glucocorticoids, indicating a dysfunctional feedback state (**Extended Data Fig. 2f, g**). It is possible that the decreased concentrations of glucocorticoids in tumor-bearing mice represent a “floor” and thus DEX cannot further decrease glucocorticoids in these mice. This would imply a potential disruption centrally or within the pituitary due to dysfunction or loss of negative feedback signaling^23,25^.

As CRH neurons within the PVN control daily oscillations in circulating glucocorticoids^18^, we examined this subpopulation of neurons more closely. We orthotopically injected EO771 tumor cells into *CRH:Ai14* mice (in which CRH cells are fluorescently labeled with TdTomato) and observed increased immunoreactivity for cFos, a canonical activity-induced transcription factor and marker of recently activated neurons, specifically in PVN^CRH^ neurons in tumor-bearing mice (**Fig. 2a-d**). This was accompanied by a small decrease in cFos expression in the SCN, which sends largely inhibitory input to PVN^CRH^ neurons to entrain circadian GC release^18^ (**Extended Data Fig. 3b-d**). These results suggest that the brain regions responsible for shaping glucocorticoid rhythms are operating differently in the presence of tumors, although a singular timepoint of analysis is not representative of how these brain regions operate over time in the context of cancer progression. Therefore, to examine hypothalamic neuronal activity with better temporal resolution during tumor progression, we used calcium fiber photometry targeting PVN^CRH^ or SCN^Vgat^ neurons (separate cohorts of mice) and monitored their 24h activity profiles (**Fig. 2e and Extended Data Fig. 3e).**

**Figure 2.**
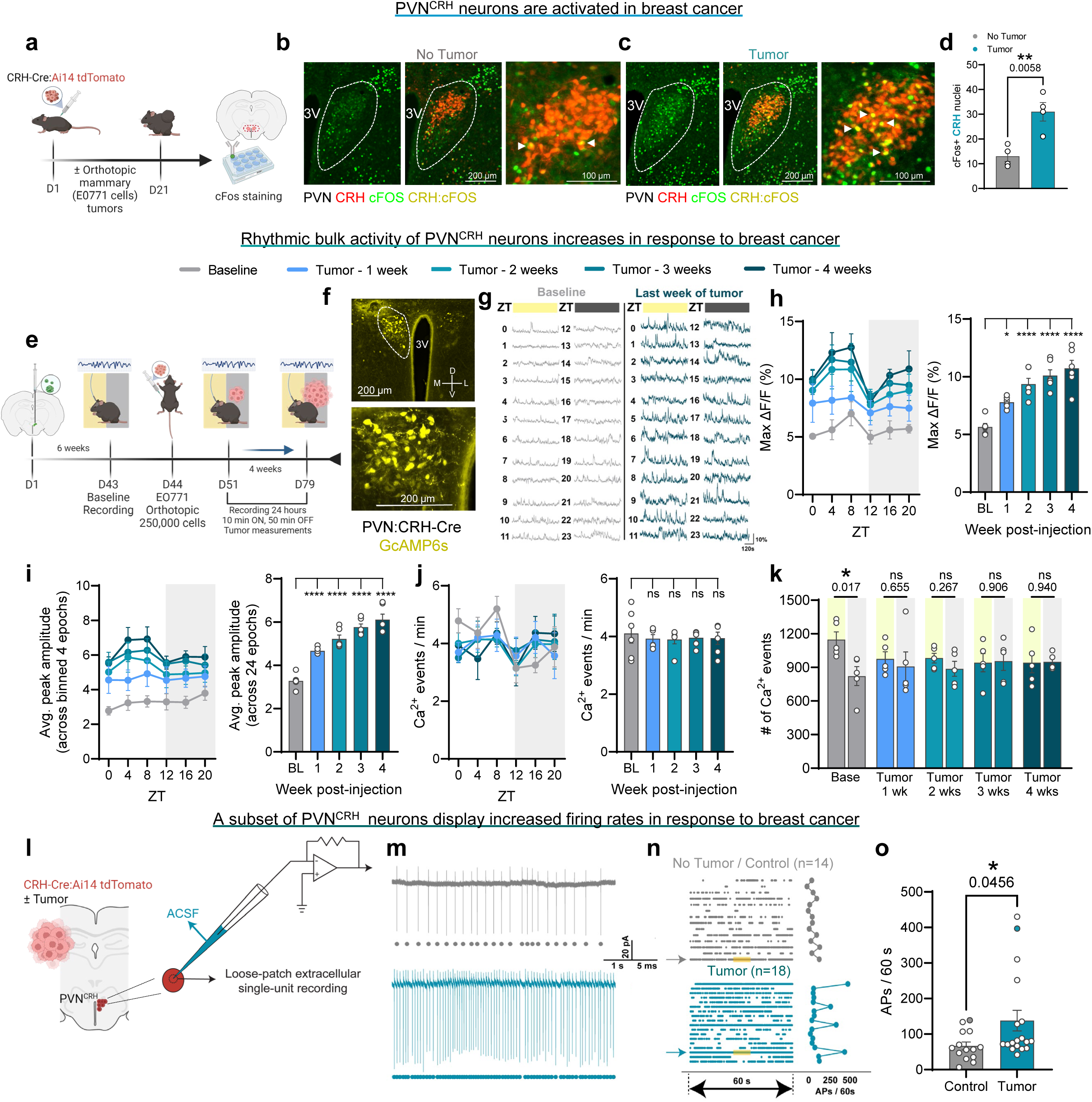
Breast tumors disrupt PVN^CRH^ neuronal activity. **a,** Timeline for cFos staining in the PVN within no tumor and tumor-bearing CRH-Cre:Ai14 mice. **b, c**, Representative coronal sections depicting cFos+ (green) nuclei and CRH (red) neurons of the PVN in control/ no tumor (**b**) and tumor (**c**) mice. **d**, Quantification of cFos+/CRH nuclei in the PVN of no tumor and tumor-bearing mice. n = 4 mice/ group. Each n is average from 2 coronal slices. **e** Cartoon depicting GCaMP6s viral injection and fiber implantation into the PVN of CRH-Cre mice, and subsequent fiber photometry recordings during tumor progression. **f,** Representative coronal section depicting GCaMP6s expression (yellow) within the PVN of a CRH-Cre+ mouse. **g,** Representative dF/F 10 min epoch traces during one day of recordings (24 epochs) from PVN^CRH^ neurons between baseline and tumor 4-week timepoints within the same mouse. (**h-j**) Photometry data for weekly recordings (10 min epochs) during tumor progression depicting max dF/F (**h**), average peak amplitude (**i**), and number of calcium transients (**j**) for PVN^CRH^ neurons collapsed into 4 hr averages (left panels) and total timepoints (right panels), respectively. n=5 mice per baseline and tumor week. **k,** Number of total calcium transients from 24-hr recordings separated by day/inactive and night/active phases. **l**, Schematic depicting loose-patch recordings of PVN^CRH^ neurons within CRH-Cre:Ai14 tumor-bearing mice. **m,** Representative current traces from PVN^CRH^ neurons in a control (upper) and tumor-bearing mouse (lower). Each dot represents an action potential fired. **n,** Raster display of population AP firing and quantification (APs fired in 60s). Portions highlighted in yellow are zoomed in panel (**m**) and are representative traces from the corresponding “filled” dots in panel **o**. **o,** Population comparison of E/I ratio between control mice (n=14, from 3 mice) and tumor-bearing mice (n=18, from 3 mice). Data represented as mean ± SEM. \**p* < 0.05, \*\**p* < 0.01, \*\*\**p* < 0.001, \*\*\*\**p* < 0.0001; two-tailed unpaired student’s t-test (**d,o**); two-way repeated-measures ANOVA followed by separate one-way ANOVAs with Bonferroni multiple comparisons (**h-k**).

### The activity of PVN^CRH^ neurons is aberrantly heightened in tumor-bearing mice

We expressed the calcium indicator GCaMP6s in PVN^CRH^ neurons and equipped mice with optic fibers to monitor bulk neuronal activity throughout the course of breast cancer progression (**Fig. 2e, f**). We collected calcium fluorescence data for 10 minutes every hour over a 24-hour period to assess circadian changes in neuronal activity (representative traces in **Fig. 2g** and schematic setup in **Extended Data Fig. 3a**). We observed a circadian rhythm in PVN^CRH^ neuronal activity, peaking slightly before the light-to-dark transition prior to tumor cell injection (i.e., baseline) (**Fig. 2h-j**). Following tumor cell injection, PVN^CRH^ neurons steadily increased their activity throughout cancer progression until the study endpoint at around 4 weeks after tumor cell injection (**Fig. 2h-j**). Increased activity was apparent one week following tumor cell injection, before tumors are palpable. Additionally, as tumors progressed, mice lost circadian rhythmicity in calcium oscillations, going from higher calcium transient frequencies during the day (inactive phase) to having similar values between the day and night (active phase) (**Fig. 2k**). The loss of circadian rhythmicity in PVN^CRH^ neurons coincides with the GC changes (**Fig. 1e**), suggesting that PVN^CRH^ neurons are a critical focal point for cancer-induced changes within the HPA axis. Notably, the elevation in neuronal activity was not observed in GABAergic neurons in the suprachiasmatic nucleus (SCN^Vgat^ neurons) (**Extended Data Fig. 3g-k**). Therefore, it is imperative to assess the electrophysiological properties of PVN^CRH^ neurons to further investigate the mechanisms underlying their aberrant activity in response to cancer progression.

We prepared acute brain slices from control and tumor-bearing *CRH:Ai14* mice to record spontaneous extracellular action potential (AP) firing from individual neurons in a loose patch configuration (**Fig. 2l**). We observed that a subset of the PVN^CRH^ neurons from tumor-bearing mice fired ∼2-3 fold more APs than most APs fired from PVN^CRH^ neurons in control animals, resulting in a significant increase of the average firing from 134 ± 22 APs / 60 s in control mice to 276 ± 58 APs / 60 s in tumor-bearing mice. (**Fig. 2m-o**). This result agrees with the observed elevation of bulk activity identified by fiber photometry. Therefore, we next examined how breast tumors distally influence synaptic input to PVN^CRH^ neurons.

### Disrupted excitatory/inhibitory balance underlies increased PVN^CRH^ neuron activity in tumor-bearing mice

We conducted whole-cell patch-clamp electrophysiology experiments to probe the mechanisms underlying increased spontaneous AP firing of PVN^CRH^ neurons in the presence of breast tumors (**Fig. 3a, b**). Increased AP firing is thought to result from changes in synaptic input (i.e., increased excitatory or decreased inhibitory input), alterations in the intrinsic properties of the neuron that changes its excitability, or a combination of the two scenarios. To assess the level of synaptic input, we recorded miniature excitatory and inhibitory post-synaptic currents (mEPSCs and mIPSCs, respectively) from PVN^CRH^ neurons in both control and tumor-bearing mice (**Fig. 3c-I**, **Extended Fig. 4a-f**). The frequency of the miniature currents is indicative of the number of synaptic vesicles released spontaneously from the pre-synaptic terminals, while the amplitude of the miniature currents reflects the sensitivity of neurotransmitter receptors on the post-synaptic PVN^CRH^ neurons. We observed a selective reduction in mIPSC frequency in tumor-bearing mice compared to controls (**Fig. 3d, i-l**), with no such changes observed for mEPSCs (**Fig. 3c, e-h**). Given that the amplitude of neither mEPSCs nor mIPSCs is affected, these results suggest that increased AP firing of PVN^CRH^ neurons could result from a reduction of pre-synaptic inhibitory input (i.e., disinhibition). Since the acquisition of mEPSCs and mIPSCs requires completely distinct bath solutions containing different pharmacological agents (e.g., antagonists for GABAergic and glycinergic receptors during mEPSCs acquisition and antagonists for AMPA and NMDA receptors during mIPSCs acquisition), we further recorded mEPSCs and mIPSCs from the same neuron to study the impact of the tumors on the relative excitatory / inhibitory input strength of single cells (**Fig. 3m-o**, and representative traces in **Extended Data Fig. 4l**). This revealed that PVN^CRH^ neurons from tumor-bearing mice are biased towards excitation, with a significantly enhanced E/I ratio (mEPSC/mIPSC frequency). This finding was further substantiated by assessment of the excitatory (Vglut2/Homer1+) and inhibitory (Vgat/Gephyrin+) synapses contacting these neurons using immunofluorescence and confocal microscopy (**Fig. 3p**). In accordance with the electrophysiological findings, we observed a significant reduction in structurally defined inhibitory synapse numbers in tumor-bearing mice with no observable changes in excitatory input (**Fig. 3q-t**). These data suggest that perturbed excitatory/inhibitory balance resulting from decreased inhibitory input may contribute to the aberrant activity of PVN^CRH^ neurons in tumor-bearing mice.

**Figure 3.**
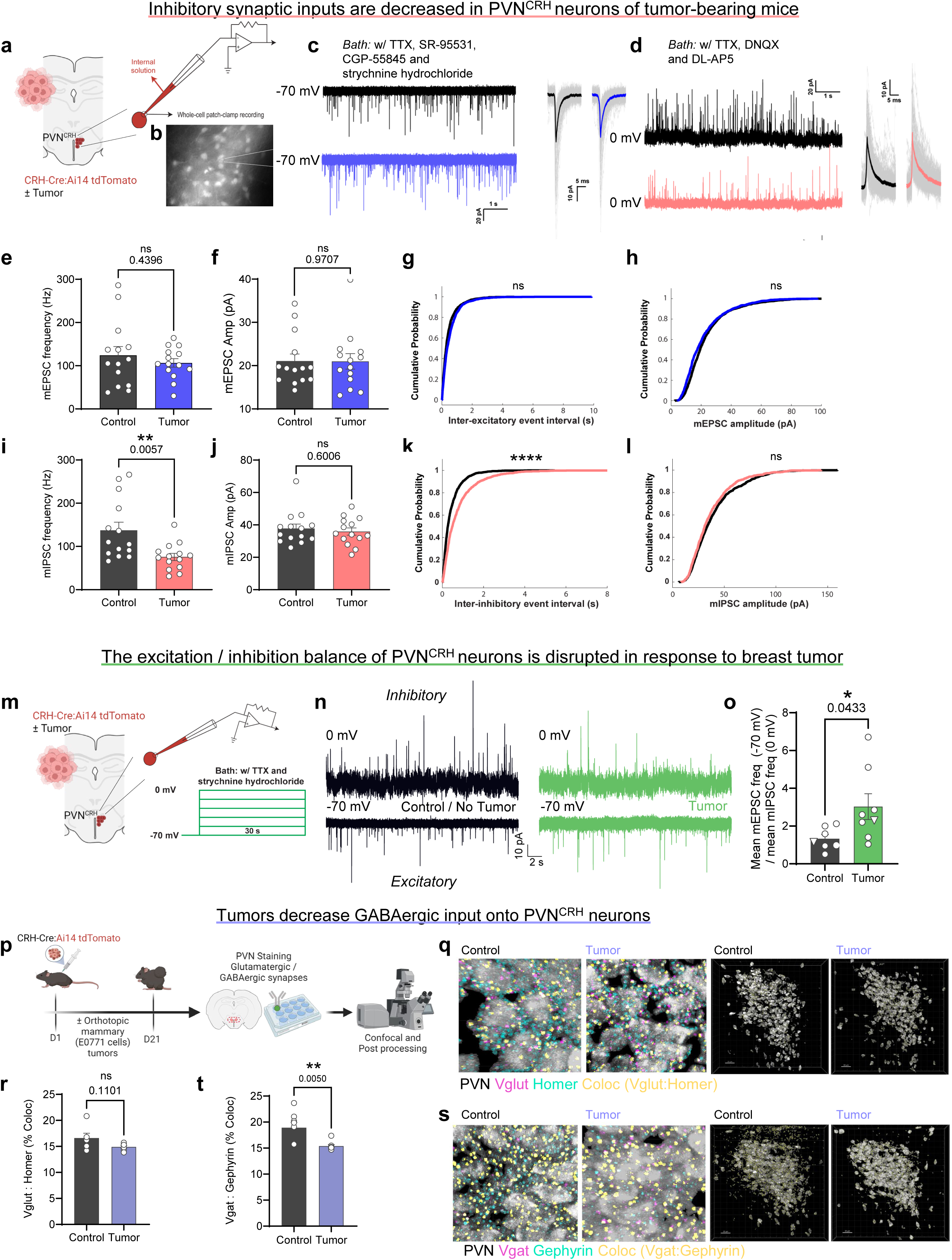
Disrupted excitatory / inhibitory input balance underlies elevated PVN^CRH^ activity. **a,** Schematic configuration of the whole-cell patch-clamp approach to record electrophysiological signals from tdtomato+ PVN^CRH^ neurons. **b,** Representative image depicting pipette patch clamp on Ai14+ CRH neurons within the PVN. **c, left:** Representative voltage-clamp traces showing mEPSCs from two PVN^CRH^ neurons clamped at −70 mV (upper: control mice; lower: tumor-bearing mice); right: each mEPSC is individually assessed (gray traces) and averaged for visual confirmation. **d,** Representative voltage-clamp traces showing mIPSCs from two PVN^CRH^ neurons clamped at 0 mV. **e-j,** Population comparison of mEPSC frequency (**e**), mEPSC amplitude (**f**), mIPSC frequency (**i**) and mIPSC amplitude (**j**) using unpaired t-test. (n=14 from 3 mice for both control and tumor for mEPSC and n=14 from 3 mice for both control and tumor for mIPSC). **g,** Population cumulative distribution frequency (CDF) of the inter-excitatory event interval, with the peak of each mEPSC trace registered as an event. **h,** Population CDF of the amplitude of mEPSCs. n=1737 for control and n=1489 for tumor. **k,** Population CDF of the inter-inhibitory event interval. The rightward shift of the CDF signifies an increase of the inter-mIPSC event interval of PVN^CRH^ neurons recorded from tumor-bearing mice, suggesting a decrease of the inhibitory neurotransmitter release. **l,** Population CDF of the amplitude of mIPSCs. n=1924 for control and n=1061 for tumor. **m**, Schematic configuration for assessing excitatory-inhibitory balance of PVN^CRH^ neurons using whole-cell patch-clamp. **n,** Representative traces showing mEPSCs and mIPSCs recorded from the same neuron at different holding voltages. Representative traces correspond to the “triangles” in panel **o**. **o,** Population comparison of the E/I ratio of PVN^CRH^ neurons between control and tumor-bearing mice. n=7 from 3 mice for control and n=8 from 3 mice for tumor. **p,** Schematic of immunostaining for Vglut, Homer, Vgat, and Gephyrin markers between tumor and non-tumor bearing CRH:Ai14 mice. **q, r,** Representative images (**q**) and puncta counts (**r**) for Vglut (pink):Homer (teal) colocalization on CRH+ neurons (white). Yellow dots represent colocalization puncta. **s, t,** Representative images (**s**) and puncta counts (**t**) for Vgat (pink):Gephyrin (teal) colocalization on CRH+ neurons (white). Yellow dots represent colocalization puncta. n=6 mice/group. Data represented as mean ± SEM. \**p* < 0.05, \*\**p* < 0.01, \*\*\*\**p* < 0.0001; two-tailed unpaired student’s t-test (**e, f, i, j, o, r, t**); Kolmogorov-Smirnov test (**g, h, k, l**).

Other than the pre-synaptic mechanism, we further investigated whether the intrinsic excitability of PVN^CRH^ neurons is changed in response to breast tumors, by injecting stepped currents to evoke APs in current-clamp mode, allowing us to extrapolate the input-output relationship of PVN^CRH^ neurons (**Extended Data 4g**). We observed that the excitability of PVN^CRH^ neurons in tumor-bearing mice is decreased (**Extended Data 4h**), without any changes in resting membrane potential (RMP, **Extended Fig. 4j**) or threshold current, which is defined as the minimum current required to evoke APs (**Extended Fig. 4k**). This result indicates that given the same input strength, PVN^CRH^ neurons in tumor-bearing mice are less likely to fire APs. This suggests that the elevated neuronal activity of PVN^CRH^ neurons is due to a pre-synaptic mechanism, where alterations in relative E/I strength overcome the reduced intrinsic excitability to yield a net bias towards increased activity.

### ‘In phase’ PVN^CRH^ stimulation attenuates primary breast tumor progression

Our work up until this point shows that breast cancer can distally influence the activity of PVN^CRH^ neurons by altering inhibitory synaptic input onto these cells (**Figs. 2, 3**). As blunted GC rhythms are associated with poor outcomes in breast cancer^1^, we tested whether re-establishing rhythmic HPA axis function influences tumor progression. We expressed the excitatory DREADD hM3Dq in PVN^CRH^ neurons (**Fig. 4b and Extended Data Fig. 6a**) and injected the ligand deschloroclozapine (DCZ^26^; 3 µg kg^-1^ IP) daily following tumor cell injection at one of two circadian timepoints (ZT 1 or ZT 10.5) (**Fig. 4a**). We reasoned that activation of PVN^CRH^ neurons at the time of day normally associated with HPA axis activation (i.e., the light-to-dark transition in mice) would re-establish glucocorticoid rhythms that resemble those observed in non-tumor bearing controls. Therefore, we injected DCZ at ZT 10.5 (1.5 hours before the light-to-dark transition), to promote ‘in-phase’ glucocorticoid release. We confirmed increases in fecal GC concentrations approximately 3 h following DCZ injections at both the ZT 10.5 and ZT 1 injection times (**Fig. 4c, d and Extended Data Fig. 5a-c, 6b, c**). We observed a significantly reduced tumor burden in mice that received ‘in phase’ (ZT 10.5), but not ‘out of phase’ (ZT 1), PVN^CRH^ neuronal stimulation (**Fig. 4e-i**), demonstrating that the anti-tumor effect of PVN^CRH^ stimulation depends on circadian phase.

**Figure 4.**
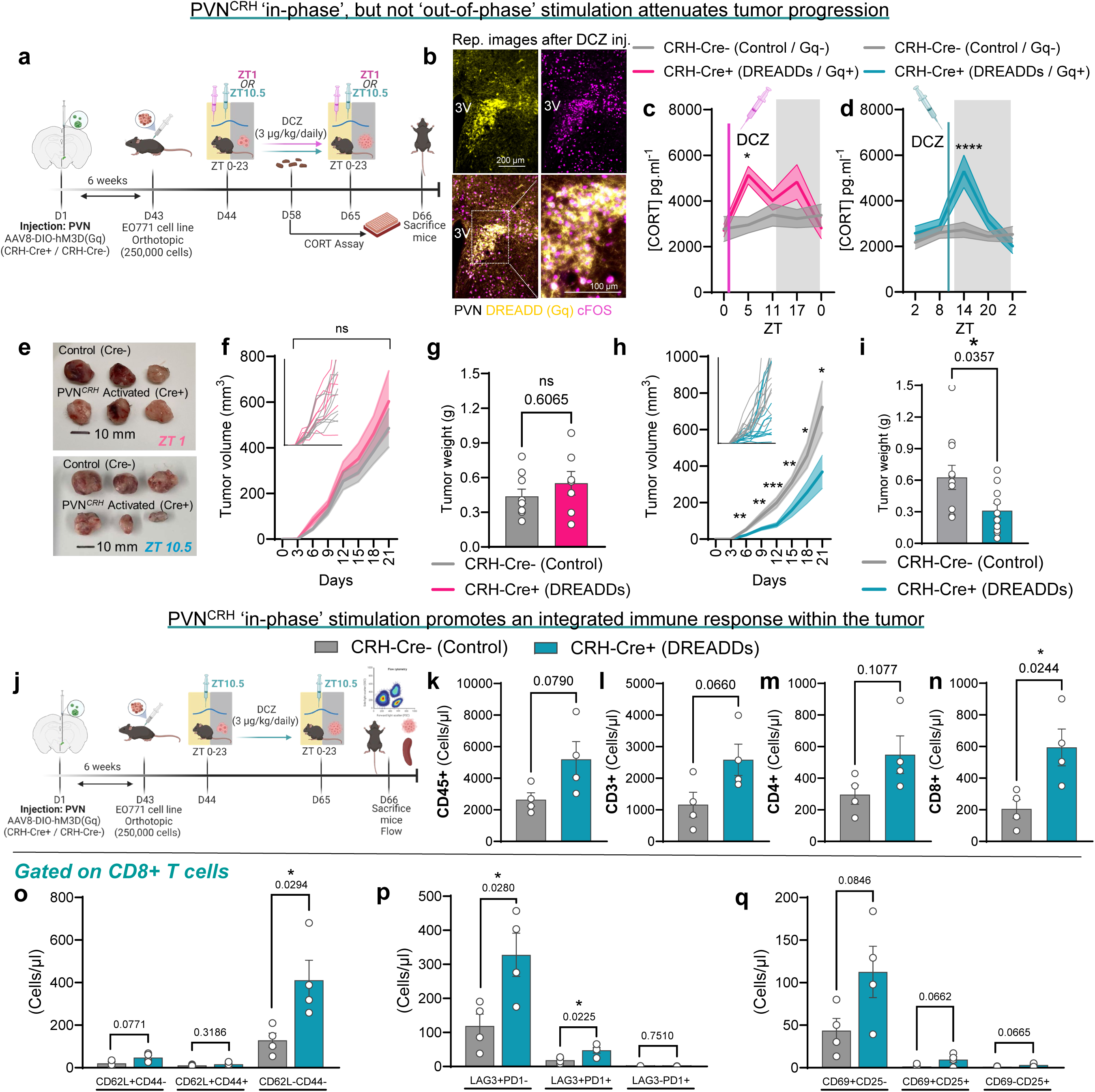
‘In-phase’ PVN^CRH^ stimulation attenuates breast tumor progression and alters the intra-tumor immune cell landscape. **a,** Schematic depicting daily, DREADD-mediated stimulation (DCZ, 3 µg/kg, ip) of PVN^CRH^ neurons at ZT1 or ZT10.5 timepoints of CRH-Cre mice during tumor progression. **b,** Representative coronal section depicting DREADD (Gq, yellow) and cFOS (purple) expression within the PVN of a CRH-Cre+ mouse after DCZ injection. **c, d,** Individual fecal corticosterone measurements across day 14 of tumor progression during a ZT1 (**c**) or ZT10.5 (**d**) DCZ injection day. n = 8-9 CRH-Cre- and 7 CRH-Cre+ mice / timepoint for ZT1; n = 10-11 CRH-Cre- and 11-12 CRH-Cre+ mice / timepoint for ZT10.5. **e,** Representative mammary tumors dissected from mice 22 days after initial tumor injection from the ZT1 (top) or ZT10.5 (bottom) DCZ-injected mice. **f, h,** Progression of mammary tumor size during 21 days after initial tumor injection from the ZT1 (**f**) or ZT10.5 (**h**) DCZ-injected mice. Inset depicts growth curves of individual mice from the mean of the larger panel. n = 9 CRH-Cre- and 7 CRH-Cre+ mice / timepoint for ZT1; n = 11 CRH-Cre- and 10 CRH-Cre+ mice / timepoint for ZT10.5. **g, i,** Final weight of EO771 tumors following sacrifice of ZT1 (**g**) or ZT10.5 (**i**) injected mice. n = 9 CRH-Cre- and 7 CRH-Cre+ mice / timepoint for ZT1; n = 11 CRH-Cre- and 10 CRH-Cre+ mice / timepoint for ZT10.5. **j,** Schematic depicting the workflow for flow cytometry of tumor immune cells after daily CRH activation in CRH-Cre-(control) and CRH-Cre+ mice. **k-n**, Quantification of CD45+ (**k**), CD3+ (**l**), CD4+ (**m**), and CD8+ (**n**) cells of CRH-Cre- and CRH-Cre+ breast tumors after daily CRH stimulation at ZT10.5 and analyzed by flow cytometry. n = 4 mice / group. **o-q**, Quantification of CD62L/CD44 (**o**), LAG3/PD1 (**p**), and CD69/CD25 (**q**) on CD8+ T cells of CRH-Cre- and CRH-Cre+ breast tumors after daily CRH stimulation at ZT10.5 and analyzed by flow cytometry. n = 4 mice / group. Data represented as mean ± SEM. \**p* < 0.05, \*\**p* < 0.01, \*\*\**p* < 0.001, \*\*\*\**p* < 0.0001; two-tailed unpaired student’s t-test (**g, i, k-q**); two-way repeated-measures ANOVA followed by separate one-way ANOVAs with Bonferroni multiple comparisons (**c, d, f, h**).

To determine whether timed GC administration also elicited a similar phenotype as top-down activation of the HPA axis (via chemogenetic stimulation of PVN^CRH^ neurons), we injected corticosterone (CORT; 10 mg kg^-1^ IP) at ZT 10.5 daily during tumor progression (**Extended Data Fig. 6d, e**). We observed no differences in tumor burden between CORT or vehicle-injected mice (**Extended Data Fig. 6f**), suggesting that enhancing activation of the HPA axis in a top-down manner is critical for influencing tumor growth. Indeed, CORT injections actively inhibited PVN activity (negative feedback; **Extended Data Fig. 6g, h**). This raises the intriguing possibility that the influence of PVN^CRH^ neurons on tumor growth is independent of downstream glucocorticoid release.

To further address that question, we asked whether timed activation of PVN^CRH^ neurons influences tumor growth via activation of tumor cell glucocorticoid receptors (GRs). Escalating concentrations of dexamethasone failed to alter proliferation in numerous breast cancer cell lines, including EO771 (**Extended Data Fig. 6l**), indicating that GR activation in the tumor cells themselves is not responsible for the change in tumor growth observed in **Fig. 4e-i**. GCs influence immune cell trafficking, but more notably, acute stimulation of PVN^CRH^ neurons alters circulating immune cell distribution^27,28^. Therefore, we assessed immune cell infiltration within tumors of stimulated mice (**Fig. 4j and Extended Data Fig. 7a**). Smaller tumors resulting from ‘in phase’ PVN^CRH^ stimulation exhibited higher numbers of immune cells (CD45+) (**Fig. 4k**). Of note, ‘in phase’ PVN^CRH^ stimulated tumors contained significantly higher numbers of CD8+ T cells (**Fig. 4l-n**) that predominantly displayed an effector phenotype (CD62L-CD44+), were highly activated (CD69+), and presented low levels of exhaustion markers (LAG3+PD1+) (**Fig. 4o-q**). We also saw significant increases in the number of NK (NK1.1+) cells and neutrophils (Ly6G+), as well as a trend towards increased B cells (CD19+) and total macrophages (Ly6C+F4/80+) (**Extended Data Fig. 7b-f**). Furthermore, bulk RNA-seq comparing the ‘in phase’ stimulated and control tumor cell population further reveals a combination effect of enhanced immune activation (e.g., *IL15ra*)^29^, elevated inflammatory responses (e.g., *Hp*, *Cd5l*)^30,31^, altered chemokine signaling (e.g., *Cxcr4*, *Ackr3*)^32,33^ and increased immunogenicity (e.g., *H60c*, *Il33*)^34,35^ (**Extended Data Fig. 6i-k**). Taken together these data indicate that ‘in phase’ PVN^CRH^ neuronal stimulation elicits an integrated anti-tumor immune response that is independent of GC release or GC action on the tumor itself. Of note, we did not observe similar immune changes in the spleens of tumor bearing mice (**Extended Data Fig. 7g**), indicating that PVN^CRH^ neuronal stimulation has tissue-specific effects.

### Tumor attenuation via PVN^CRH^ neuronal stimulation requires CD8+ effector T cells

To determine whether the attenuated tumor growth phenotype is dependent on this integrated immune response, we treated mice with depleting anti-CD8α antibodies (or IgG2 isotype control antibodies) every 4 days in the context of daily Gq DREADD stimulation of PVN^CRH^ neurons (**Fig. 5a**). We hypothesized that the anti-tumor response due to ‘in-phase’ PVN^CRH^ stimulation is, at least in part, due to CD8+ T cell infiltration into the tumor. Treatment of mice with anti-CD8α antibodies depleted CD8+ T cells within the tumor (without affecting CD4+ T cells) (**Fig. 5e, f and Extended Data Fig. 7h**), in addition to abolishing the anti-tumor effect of ‘in-phase’ PVN^CRH^ stimulation (**Fig. 5b-d**). This result shows that CD8+ T cells play a key and non-redundant role in transducing the neuronal signal into an anti-tumor immune response.

**Figure 5.**
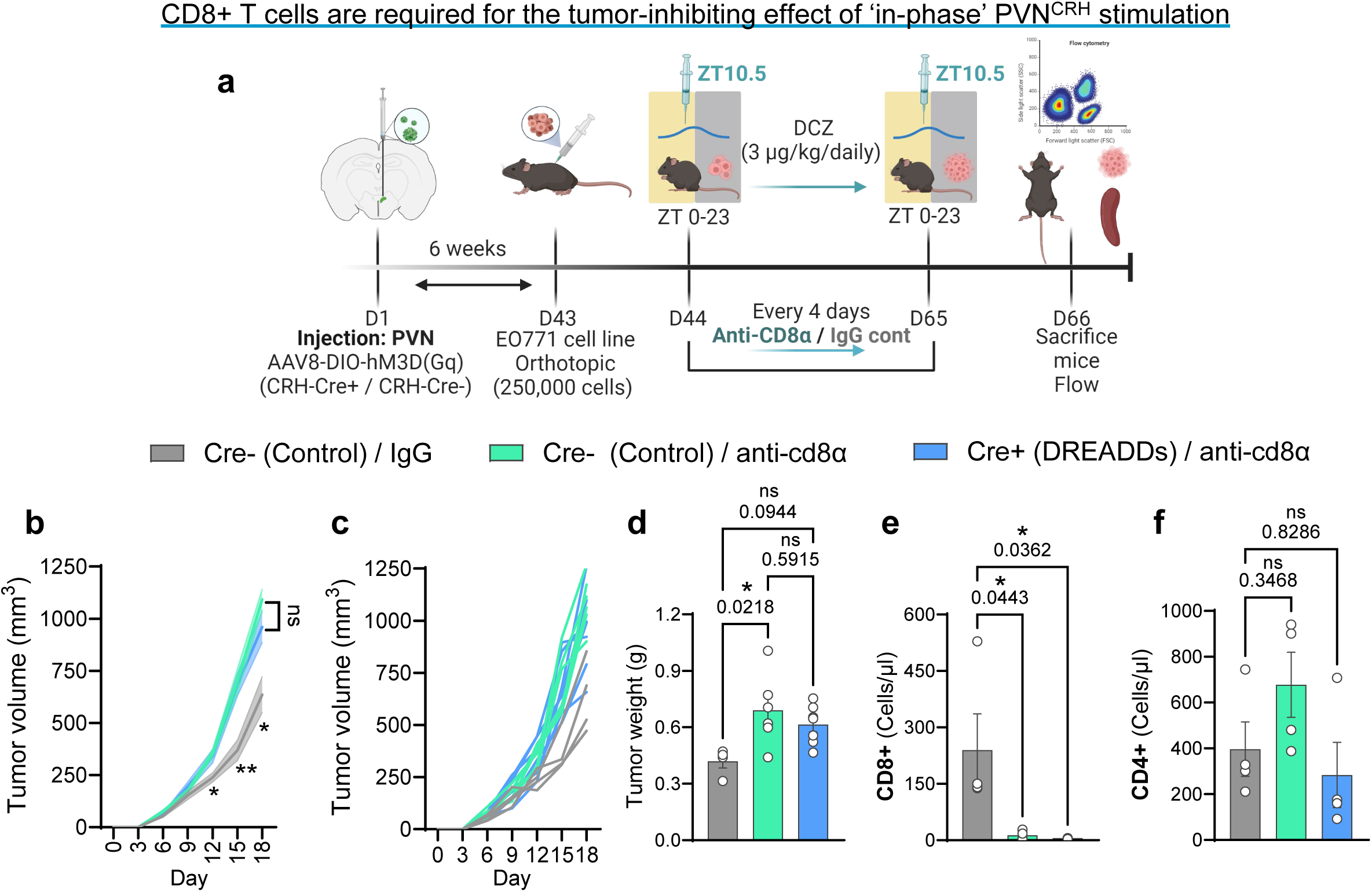
CD8+ T cells drive the tumor-inhibiting effect of ‘in-phase’ PVN^CRH^ stimulation. **a,** Schematic depicting the workflow for flow cytometry of tumor immune cells after daily CRH activation in control and CD8+ depleted mice. **b, c,** Progression of mammary tumor size in ZT10.5 DCZ-injected mice during IgG control or CD8+ depletion. n = 4 Cre-/control; n = 6 Cre-/anti-CD8α; n = 7 Cre-/anti-CD8α. **d,** Final weight of mammary tumors in ZT10.5 DCZ-injected mice after IgG control or CD8+ depletion. n = 4 Cre-/control; n = 6 Cre-\anti-CD8α; n = 7 Cre-\anti-CD8α **e, f,** Quantification of CD8+ (**e**) and CD4+ (**f**) breast tumor cells in ZT10.5 DCZ-injected mice after IgG control or CD8+ depletion and analyzed by flow cytometry. n = 4 mice / group. Data represented as mean ± SEM. \**p* < 0.05, \*\**p* < 0.01; two-way repeated-measures ANOVA followed by separate one-way ANOVAs with Bonferroni multiple comparisons (**b**); one-way ANOVAs with Bonferroni multiple comparisons (**d-f**).

## Discussion

Circadian rhythms in glucocorticoid concentrations play a major role in maintaining systemic health^17^. Identifying the causal factors driving aberrant GC circadian rhythms in people with breast cancer has remained challenging in part due to the many co-variates that can each independently influence GC concentrations (e.g., age, cancer treatment regimen, diet). Cancer cells modify their local microenvironment to ensure an adequate supply of metabolic fuel and to evade immune destruction. Less well appreciated, however, is the ability of cancers to influence the activity of distal host tissues^36^. Here, we demonstrate that breast cancer itself can disrupt host circadian GC rhythms via distal disinhibition of discrete hypothalamic neurons (i.e., PVN^CRH^ neurons). We reveal that this normal diurnal variation has anti-tumor effects: when we reestablish robust GC rhythms via chemogenetic activation of PVN^CRH^ neurons, we can attenuate tumor growth, an effect not observed when these neurons are stimulated ‘out-of-phase’ with the natural circadian cycle. Importantly, this reduction in tumor growth relies not on GC signaling but on the actions of CD8+ T cells, as tumors become unresponsive to PVN^CRH^ ‘in-phase’ stimulation in the absence of these effector cells.

Our results raise three major questions: (1) what is the signal(s) from the tumor-bearing host that distally influences PVN^CRH^ neuron function?; (2) What cells are upstream from PVN^CRH^ neurons that are responsible for reduced inhibitory input?; and (3) How does ‘in-phase’, but not ‘out of phase’ PVN^CRH^ neuronal stimulation promote an integrated anti-tumor immune response?

We detect changes in hypothalamic neuronal activity as soon as one week following tumor cell injection (**Fig. 2**), so it is implausible that a neuronal circuit between PVN^CRH^ neurons and the mammary tumor can form and then drive the response given the time constraints on axon outgrowth within adult animals^37^. Polysynaptic connections between the (healthy) mammary gland and PVN have been mapped in rats^38,39^. Of these neurons, oxytocinergic neurons (PVN^OXT^) that facilitate milk letdown during lactation are likely a key population^40^. Interestingly, PVN^OXT^ neurons suppress dextran sodium sulfate (DSS)-induced colorectal tumor growth via inhibition of downstream sympathetic output^41^, suggesting that multiple hypothalamic neuronal populations can act in concert to influence cancer. Alternatively, a humoral signal may relay information from the periphery to the brain to activate PVN^CRH^ neurons early in breast cancer development. Likely candidates for this are pro-inflammatory cytokines, which play a major role in activating the HPA axis to putatively prevent runaway inflammation in the face of acute immune challenges^42–44^. During inflammation, PVN^CRH^ neurons rapidly upregulate their expression of interleukin-6 receptors (IL-6Rα)^42^, and injections of interleukin-1 induce a delayed and long-lasting (at least 3 weeks) increase of vasopressin stores in CRH axon terminals leading to hyperresponsiveness of the HPA axis to subsequent stimuli^44,45^. Additionally, prostaglandin E2 (PGE_2_) reduces GABAergic input to paraventricular neuroendocrine cells by binding presynaptic EP3 receptors to drive enhanced HPA-axis output^46^. It will be important to determine if CRH neurons are directly sensitive to tumor-induced changes in systemic physiology, or whether they become activated by an initial receiver cell (e.g., brain endothelial cell, astrocyte, microglia) which then relays the signal. Recent work in a mouse model of colorectal cancer showed that tumor derived interleukin-6 travels through the circulation and binds IL-6Rα directly on Gfral-expressing neurons in the area postrema, where it enhanced neuronal excitability to suppress food intake and promote cancer cachexia^47^. Data such as these suggest that peripheral cytokine-to-brain signaling may underlie multiple aspects of systemic disruption in cancer. Parabiosis or plasma-transfer experiments will help in determining whether tumors activate PVN^CRH^ neurons via a neural or humoral route.

We show that inhibitory input to PVN^CRH^ neurons is reduced in tumor-bearing mice (**Fig. 3**), suggesting that the critical population influenced by breast cancer is upstream from these cells. We suspect that reduced input from GABAergic neurons within the SCN, which normally act to entrain PVN^CRH^ neurons^18^, drives this disinhibition. Several other GABAergic populations innervate PVN^CRH^ neurons, including those in the lateral hypothalamic area, dorsomedial hypothalamus (DMH), ventrolateral preoptic area, as well as extra-hypothalamic areas like the bed nuclei of the stria terminalis (BNST)^48^. Of these, the DMH plays a major role in transducing circadian information to the HPA axis, as it also receives direct input from the SCN, and lesions to the DMH reduce circadian glucocorticoid rhythms by up to ∼89%^49^. We show reduced cFos expression within the SCN of tumor bearing mice (**Extended Data Fig. 3**), but retrograde tract tracing studies will be essential in confirming that reduced input from these cells drives aberrant PVN^CRH^ activity in response to breast cancer.

Regarding the effect of PVN^CRH^ neuron stimulation on tumor growth, our results contrast with recent work showing that chemogenetic activation of PVN^CRH^ neurons promotes the growth of subcutaneously implanted Lewis lung carcinoma (LLC) cells, an effect associated with reduced infiltration of CD8+ T cells into the tumor^50^. Unfortunately, the time of day that neurons were stimulated during tumor growth is not reported in that paper, making comparison to our dataset difficult. We find that the effect PVN^CRH^ neurons have on tumor growth depends on circadian phase, and future studies should carefully consider time-of-day as a critical biological variable^51^. This dependence may be driven by differential sensitivity to ACTH within the adrenal gland across the day^19^, or differences in the structure of the glucocorticoid rhythm induced by PVN^CRH^ stimulation, which differ between mice stimulated at ZT1 vs. ZT10.5 (**Fig. 4**). Acute stress or activation of PVN^CRH^ neurons promotes the redistribution of immune cells from circulation into the bone marrow and lymphoid tissues^27,28^. Our data suggest that activation of these neurons also promotes the infiltration of effector CD8+ T cells into tumor tissue, but this effect is powerfully modulated by circadian phase (**Fig. 4 and 5**). This may be due to the actions of glucocorticoids on chemokine receptor expression in immune cells^52,53^, rendering them more sensitive to chemoattractant cues at different times of the day in line with the body’s physiological needs. Indeed, glucocorticoids drive daily oscillations in T cell IL-7R expression which induces differential distribution of T cells to lymph nodes, spleen, and blood via modulation of CXCR4 expression^14^. This is important to consider as glucocorticoids are widely thought to primarily promote cancer growth and metastasis^54^, whereas our data suggest that their actions on tumor growth and anti-tumor immunity can vary substantially according to the circadian cycle^53^.

Multiple subpopulations of PVN^CRH^ neurons exist^55^, with two main categories being parvocellular neurosecretory and non-neurosecretory cells (PNCs and PNNCs, respectively). PNC CRH neurons produce CRH and release it into the hypophyseal portal system at the median eminence to act in the canonical HPA axis, while PNNC CRH cells are primarily centrally projecting pre-autonomic neurons that control sympathetic outflow to the periphery by innervating multiple autonomic nuclei^56^. This raises the question: which sub-population(s) of CRH neurons are responsible for influencing tumor growth? Our electrophysiology data suggests that a subset of the total CRH population drives the change in E/I balance caused by tumors (**Fig. 2 and 3**). We reasoned that if PNC CRH neurons are responsible for the anti-tumor effect of PVN^CRH^ neuron stimulation, then timed daily injections of exogenous corticosterone ‘in-phase’ with the natural rhythm would confer benefits in mouse models of breast cancer. Our data, however, indicate that ‘in-phase’ corticosterone administration slightly *increases* tumor progression (**Extended Data Fig. 6**), and this is associated with a reduction in cFos immunoreactivity within PVN^CRH^ neurons. This suggests that activation of PVN^CRH^ neurons influences cancer progression independently from the actions of glucocorticoids, indicating that PNNC pre-autonomic CRH neurons are likely the critical mediators of their effects on tumor growth. Projection-specific manipulations and single-cell RNA-sequencing will be key in answering this question.

Neuromodulation of PVN^CRH^ neurons represents a new strategy for combating cancer and, importantly, represents a completely orthogonal and non-redundant approach to existing therapies (e.g., chemotherapy). Clearly, much more work is required to capitalize on these discoveries, but our work provides prima facie evidence that breast cancer disrupts normal brain-body feedback loops (i.e., HPA axis) via distally disinhibiting PVN^CRH^ neurons, and that enforcing host circadian rhythms via timed neuromodulation may offer a novel therapeutic strategy towards eliminating breast cancer. As low frequency stimulation of vagal (cranial nerve X) afferents activates the HPA-axis^57^, emerging bioelectronic medicine approaches that target the vagus nerve^58^ may comprise feasible strategies to modulate endogenous HPA-axis circadian rhythms in patients with breast cancer.

## Methods

### Animals

All experimental procedures and protocols were approved by the Institutional Animal Care and Use Committee of Cold Spring Harbor Laboratory (CSHL) and performed in accordance with the US National Institutes of Health (NIH) guidelines. Female mice (∼3-6 months of age; ∼20-28 g) were group housed, with the exception of mice undergoing fiber photometry who were single housed to protect integrity of the fiber optic, under 12 hr:12 hr light:dark cycle (lights on at 7:00 AM) within a room with temperatures (∼22 °C) and humidity ∼45%. Food (LabDiet, #LD505330LB) and water were provided ad libitum. All mice were bred on a c57Bl6 background, and subsequent breeding took place at Cold Spring Harbor Laboratory. All breeding pairs were originally purchased from The Jackson Laboratory, consisting of: C57BL/6J (Wildtype/WT, #000664), B6(Cg)-*Crh^tm^*^1^*^(cre)zjh^/*J (CRH-ires-CRE, #01204), *Slc32a1^tm^*^2^*^(cre)low^l*J (Vgat-ires-CRE, #016962) and B6.Cg-*Gt(ROSA)26Sor^tm^*^14^*^(CAG-tdTomato)Hze^*/J (Ai14, #007914), B6.FVB-TG(MMTV-PyVT) 634Mul/LellJ (MMTV-PyMT, #022974). All mice were acclimated to the housing conditions at least 1 week prior to any manipulation (surgery, CORT measurements, etc.).

### Drugs & in vivo depleting antibodies

The following drugs were utilized in the experiments: CORT (Corticosterone HBC Complex, 10 mg/kg i.p., Sigma-Aldrich, #C174, injected at ZT 10.5); ACTH (ACTH 1-39 Porcine, 5 and 50 mg/kg i.p., BOC Sciences, #B2693-194638, injected at ZT 0); DEX (Dexamethasone, 10 mg/kg i.p., Sigma-Aldrich, #D4902, injected at ZT 1); DCZ (Deschloroclozapine dihydrochloride – water soluble, 3 µg/kg i.p., Hello Bio, #HB1126, injected at ZT 1 and ZT 10.5); IgG Control (*InVivo*MAB rat IgG2b isotype control, BioXcell, #BE0090, injected every 4 days at ZT 4); and Anti-CD8 (InVivoMAb anti-mouse CD8α, 100µg/mouse, BioXcell, #BE0061, injected every 4 days at ZT 4).

### Stereotaxic Surgery (Viral Injections and Optic Fiber implants)

All mice surgical procedures were in line with CSHL guidelines for aseptic technique. Mice were anesthetized with isoflurane (3% induction/1.5% maintenance; Somnosuite, Kent Scientific), and then injected with buprenorphine SR (Zoopharm, 0.5 mg/kg, s.c.). Upon confirmation of deep anesthesia mice were placed into a stereotaxic frame (David Kopf Instruments) where they were maintained at 1-1.5% isoflurane. A midline incision was then made from the posterior margin of the eyes to the scapulae to expose the braincase. The skull was cleaned and then a drill was be positioned over the skull to drill a hole for the viral injection. Mice were then injected unilaterally within the PVN (−0.85 mm AP, ±0.2 mm ML, −4.7 mm DV) or SCN (−0.5 mm AP, ± 0.15 mm ML, −5.6 mm DV) using a 30-gauge blunt Neuros syringe (Hamilton) at a rate of 50 nl/min for a total volume of 0.3microL. All viruses were obtained from Addgene, and consisted of AAV9-CAG-FLEX-GCaMP6s (viral titer 2 x 10^13^); AAV8.hSyn-DIO-HA-hM3D(Gq)-IRES-mCitrine (viral titer 2 x 10^13^); AAV8.hSyn-DIO-hM3D(Gq)-mCherry (viral titer 2.6 x 10^13^) AAV5-EF1α-DIO-hChR2-(H134R)-eYFP (viral titer 1.8 x 10^13^). After the infusion, the needle was left in place for at least 10 mins before the microinjector (World Precision Instruments) was withdrawn slowly. Directly following virus injection, a fiber optic cannula (400 μm in diameter; 0.48 NA, Doric Lenses) was lowered just dorsally to the injection site. The optical implant was then cemented in place with Metabond (Parkell) and dental cement. After surgery, mice were then allowed to recover until ambulatory on a heated pad, then returned to their home cage with Hydrogel and DietGel available. Mice were then allowed to recover for ∼4-5 weeks to allow for viral expression before behavioral experiments and fiber photometry recordings began. At the end of experiments viral expression within the specified brain regions were validated via visualization of fluorescence expression.

### Fiber photometry

Approximately ∼4-5 weeks after viral injection and fiber optic implantation, mice began baseline recording sessions. In brief, mice were tethered to a fiber optic cable, within their home cage, via a ceramic mating sleeve connected to the implanted optic fiber, and fiber photometry data was collected using a fiber photometry setup with optical components from Doric Lenses controlled by a real-time processor from Tucker Davis Technologies (TDT; RZ10X). TDT Synapse software was used for data acquisition. 465 nm (Signal / GCaMP) and 405 nm (Control / Isosbestic) LEDs are modulated at 211 or 230 Hz and 330 Hz, respectively. LED currents were adjusted in order to return a voltage between 100 and 150 mV for each signal, and were offset by 5 mA. The signals were then demodulated using a 6 Hz low-pass frequency filter, where subsequent analysis took place. Fiber photometry data analysis took place utilizing Guided Photometry Analysis in Python (GuPPy)^59^.

### Cell culture and injections

E0771 mammary cells (i#CRL-3461; ATCC) obtained from the supplier were cultured in Dulbecco’s modified Eagle’s medium supplemented with 10% <u>fetal bovine serum</u> and 1% penicillin/streptomycin. Cells were used within 3 passages upon arrival. These cells (EO771), when implanted, display a breast cancer phenotype similar to the basal subtype^20^. Cells were incubated at 37°C in a mixture of 5% carbon dioxide and 95% air. Once cells reached ∼90% confluence, cell numbers and viability were determined with a 1:1 mixture of trypan blue (Life Technologies) in an automated hemocytometer (The Countess II, Life Technologies, Carlsbad, CA, USA). Cells were diluted in DMEM to make a final concentration of 2.5 x 10^5^ cells per 100 μL. Prior to injection, EO771 cells (approximately 80-90% confluence) were trypsin detached, filtered to prevent cell clumping, mixed with a 1:1 growth factor–reduced Matrigel™ Matrix (BD Matrigel™ Matrix, BD Biosciences, Bedford, MA): PBS mixture and kept on ice until administration. Mice were briefly anesthetized under isoflurane, and received subcutaneous injections (100 μL surrounding the right inguinal mammary fat pad) of E0771 in DMEM (hereafter referred to as “Tumor”) or DMEM alone (“No Tumor/Sham”). Proper injection was confirmed by observing the presence of a wheal upon injection. Mice were inspected approximately every 3 days for tumor growth. Once a tumor became palpable, measures were taken daily using sliding calipers by the same person and volume was determined using the formula Volume = (length x width^2^)/2^60^. If any mouse displayed signs of tumor ulceration, the mouse was taken out of the study and sacrificed accordingly.

### Fecal Corticosterone Assay

Mice were placed in a clean cage every 6 hrs for approximately 10 mins, where 3-4 fecal pellets were collected, placed in sterile Eppendorf tubes, and stored at −80 °C until processing. If mice did not defecate after 5 mins of placement in a clean cage, the mice were returned back to their home cage. Samples were then placed into an oven at 58°C and were allowed to dry completely (∼3-4 hours) until the fecal pellets no longer decreased in weight. The samples were then placed in a TissueLyser II (Qiagen) and ground into a fine powder. Samples were then weighed, and 100% ethanol was added so that the final concentration of the fecal sample yield equaled 100 mg/ml. Samples were then allowed to rotate overnight at room temperature. Samples were then centrifuged 15 mins, 5000 x rpm, at 4°C, and the ethanol supernatant was collected. Samples were then prepared and diluted based on corticosterone enzyme immunoassay kit protocol (Arbor Assays, #K014-H1) so that the total ethanol content <5%. Samples were further processed in duplicate for corticosterone concentration (Corticosterone ELISA Kit, Arbor Assays), where the intra-assay coefficients of variation were <10%, and the optical density of each sample was assessed by a microplate reader at 450 nm (Spectramax I3x, Molecular Devices).

### RNA Extraction and Quantitative RT-PCR

Following euthanasia via Euthasol (Virbac), tissues were immediately dissected and placed on ice. RNA was extracted using TRl reagent (Sigma-Aldrich) according to the manufacturer’s instructions, precipitated with isopropanol and purified with 75% ethanol. RNA quality and quantity were determined using a spectrophotometer (NanoDrop), and cDNA was synthesized using MMLV reverse transcription (Lucigen). 20 ng of cDNA/reaction was used in subsequent PCR. Taqman Fast advanced master mix (Life Technologies) containing AmpliTaq Fast DNA polymerase was used in a 10 ul triplex reaction with the Taqman target gene/probes. The primer/probe pairs consisted of: *Cyp11b,1* Mm01204952_m1; *Hsd3b1,* Mm01261921_mH; *StAR,* Mm0441558_m1; *Mc2r,* Mm00434865_s1; *Pomc,* Mm00435874_m1; *NR3c1,* Mm00433832_m1. The 2-step real-time PCR cycling conditions used were: 95 °C for 20s, 40 cycles of 95 °C for 1s, and then 60 °C for 20s. Relative gene expression was calculated using the 2^(-delta-delta Ct) method and normalized to 18s rRNA.

### Bulk RNA-Seq and data analysis

Total RNAs were extracted from CRH-Cre (+/-) mice tumors with DCZ-treatment using Trizol reagent and purified with isopropanol and ethanol. cDNA libraries were prepared using the Kapa mRNA HyperPrep kit for Illumina® Platforms (Roche, Cat. 08098115702) following the manufacturer’s protocol. Sequencing was performed using the NextSeq 2000 P1 300 cycle kit (Illumina) to achieve a target output of 200 million paired-end reads. Bulk RNA-seq data were analyzed using a JupyterHub-based pipeline. Initial quality control of the raw sequencing data was performed with FastQC. Reads were aligned to the mouse reference genome GRCm39 using STAR, followed by gene quantification with featureCounts. Differential expression analysis was conducted using DESeq2, with the False Discovery Rate (FDR) threshold of <0.05.

### Acute slice electrophysiology

CRH-Cre x Ai14 mice were anesthetized by inhaling isoflurane, rapidly decapitated and the brain slices containing the paraventricular nucleus of the hypothalamus (approximate AP coordinates −0.75mm to −0.95mm) were removed and submerged in oxygenated artificial cerebrospinal fluid (aCSF, in mM: 125 NaCl, 2.5 KCl, 25 glucose, 1.25 NaH_2_PO_4_, 25 NaHCO_3_, 2 CaCl_2_, 1 MgCl_2_, 0.4 Ascorbic acid, 3 Myo-inositol and 2 Pyruvic acid, osmolarity adjusted to 300–305 mOsm, and saturated with 95% O_2_/ 5% CO_2_) at RT. The brain was glued (Loctite 404, Henkel, <u>Düsseldorf, Germany</u>) to the stage of a vibrating microtome (7000smz-2, Campden instrument, Loughborough, England). Coronal sections (200-300 μm) containing CRH+ cells were collected, submerged in an incubation chamber (model 7470, Campden Instrument, Loughborough, England) of continuously oxygenated aCSF, and incubated at 37°C for 1 h. Slices were then transferred to a recording chamber and submerged in aCSF. The chamber solution was held at 34 °C through an inline feedback temperature controller (TC-344C, Warner Instruments, Hamden, CT, USA). CRH+ cells were identified under fluorescent illumination (pE-400 Max, CoolLed, Hamshire, UK) and examined using an infrared-differential interference contrast optics microscope (SliceScope, Scientifica, Uckfield, UK). The recording chamber was continuously perfused with aCSF at a rate of 2– 4 ml/min.

Borosilicate capillary glass pipettes (1B120F-4, WPI, Sarasota, FL, USA) were pulled to a resistance of 4–8 MΩ with a two-stage puller (PC-100, Narishige, Tokyo, Japan) and back-filled with internal solution. The internal solution contained the following in mM: 145 K-gluconate, 5 KCl, 5 EGTA, 10 HEPES, 1 MgCl_2_, 4 Na_2_ATP, and 0.3 Na_2_GTP, pH 7.2 adjusted with KOH and osmolarity adjusted to 295 mOsm. In voltage-clamp for loose-patch experiments, bath aCSF was used as internal solution. The holding voltage was set to 0 mV during data acquisition (D. Anson and M. Roberts, 2002). In voltage-clamp for post-synaptic current recordings, cesium-based internal solution (125 CsMeSO_4_, 15 CsCl, 5 EGTA, 10 HEPES, 1 MgCl_2_, 4 Na_2_ATP, and 0.3 Na_2_GTP) was used. 5 mM QX314 was added to the internal solution to prevent antidromic action potentials, and 1 μM TTX was added to bath solution. For mEPSC recordings, SR-95531 (5 μM), CGP-55845 (2 μM) and strychnine hydrochloride (1 μM) were added to bath solution to block GABAergic / glycinergic receptors. For mIPSC recordings, 6,7-Dinitroquinoxaline-2,3-dione (DNQX, 40 μM) and DL-2-Amino-5-phosphonopentanoic acid (DL-AP5, 50 μM) were added in bath solution to block the AMPA and NMDA receptors, respectively. For E/I ratio experiments where both mEPSC and mIPSC were collected from the same neuron, only TTX and strychnine hydrochloride were added to the bath solution. In current-clamp, a liquid junction potential from 7-10 mV was corrected for potassium-based internal solutions. All electrophysiological signals were amplified, series resistance-compensated at 60% to 80%, and low-pass filtered at 10 kHz (MultiClamp 700B, Axon Instruments, Novato, CA, USA), digitized at 20 kHz (lab-written customized Matlab routines controlling a USB-6363 DAQ board from National Instrument, Austin, TX, USA), and stored for offline analysis in Matlab (Mathworks, R2023, Natick, MA, USA).

### Confocal microscopy and quantification of synapses

Images were acquired using a Zeiss LSM710 laser scanning confocal microscopy with the Plan-Apochromat 40x/1.3 oil immersion objective. Z-stacks of 40 μm were applied for each brain slice in a 3D context, comprising 40 slices with z-stack intervals of 1 μm each. Imaris software (10.1.1, Oxford Instrument) was used for precise quantification of pre-/post-synaptic 3D object-based colocalization within the confines of the CRH soma area. The spot function and background correction functions in Imaris were applied to measure the fluorescence intensity and centroid positions of Homer, Vglut2, Gephyrin and Vgat signals in each image stack. The number of colocalized voxels and the percentages of each synaptic volume above the colocalized threshold were quantified. Uniform imaging and analysis parameters were applied to both control and tumor samples to ensure methodological consistency.

### Immunohistochemistry

Mice were transcardially perfused with 0.1 M phosphate-buffered saline (PBS) and then 40 mL 4% paraformaldehyde (PFA). For colocalization experiments (**Fig. 3p-t**), mice were perfused with a glyoxal fixative consisting of a PBS mixture of 9% glyoxal and 8% acetic acid^61^. Brains were dissected and post-fixed in 4% PFA overnight and then transferred to 30% sucrose solution for cryoprotection. Brains were then sectioned at 35 μM on a cryostat (Leica CM1950) and stored in a 0.1M PBS at 4°C prior to immunohistochemistry experiments. For immunohistochemistry free-floating sections were washed in 0.1 M PBS for 3 × 10 minutes intervals. Sections were then placed in blocking buffer (0.1% Triton X-100 and 3% bovine serum albumin in 0.1 M PBS) for 1 hr at room temperature. After blocking buffer, sections were placed in primary antibody (cFos, 1:2000, Cell Signaling, #2250; Vglut, 1:500, Invitrogen, #PA5-119621; Vgat, 1:500, Synaptic Systems, #131-004;-Homer, 1:500, Synaptic Systems, #160-026, Gephyrin, 1:500, Synaptic Systems, #147-011) for approximately 16-20 hrs at room temperature. After 3 × 10-minute 0.1 M PBS washes, sections were incubated in secondary antibody (AlexaFluor 488 goat anti-mouse, 1:1000, Life technologies, #A21121; AlexaFluor 488 goat anti-chicken, 1:1000, Life technologies, #A32931; AlexaFluor 647 goat anti-rabbit, 1:1000, Life technologies, #A21245; AlexaFluor 647 goat anti-guinea pig, 1:1000, Life technologies, #A21450) for 2 hours at room temperature, followed by subsequent washes (3 X 10 minute in 0.1 M PBS). Sections were then mounted and coverslipped with Vectashield mounting medium with DAPI (Vector Laboratories, H-2000) and imaged on a Keyence BZ-X800 and Leica TCS SPE confocal microscope. For cohorts, viral expression and optical fiber placements were confirmed before inclusion in the presented datasets.

### Single cell suspensions & Flow cytometry

Breast tumors were harvested 3 weeks post-injection and processed for flow cytometry analysis. Tumors were first manually minced to pieces <1 mm and incubated for 30 min at 37°C with vigorous shaking in digestion solution containing 1x collagenase/hyaluronidase (Stem Cell # 07912), 20U/mL DNase 1 in 5 mL RPMI 1640 Glutamax medium (ThermoFisher #61870036) with 5% FBS. Then cell suspensions were centrifuged at 1275 rpm for 5 minutes at 4°C, washed with HBSS + 5% FBS, and filtered through a 100μm cell strainer (Falcon). The resulting suspensions were centrifuged again at 1275 rpm for 5 minutes at 4°C and resuspended in 2 mL ACK lysis buffer to eliminate blood cells. Cells were then washed with PBS and resuspended in FACS buffer and used for immediate analysis or fixed with Fixation Buffer (BD Cytofix, BD Biosciences, 554655, 3199180). The following fluorophore-conjugated and their indicated dilutions consisted of: For Panel 1: CD45.2 BV711 (clone 104, BioLegend, 109847, B348415, 1:100 dilution), CD3 PE/Cyanine7 (clone 17A2, BioLegend, 100220, B401339, 1:100 dilution), CD19 BV650 (clone 1D3, BD Biosciences, 563235, 3347372, 1:100 dilution), CD11c BV785 (clone N418, BioLegend, 117335, B295112, 1:100 dilution), CD11b BUV395 (clone M1/70, BD Biosciences, 563553, 3346840, 1:100 dilution), Ly-6C APC-Cyanine7 (clone HK1.4, BioLegend, 128026, B309226, 1:100 dilution), Ly-6G BV605 (clone 1AB, BD Biosciences, 563005, 3187156, 1:100 dilution), F4/80 BV421 (clone BM8, BioLegend, 123131, B367651, 1:100 dilution), NK1.1 PerCP/Cyanine5.5 (clone PK136, Tonbo Biosciences, 65-5941-U100, C5941021417653, 1:100 dilution), MHC-II FITC (clone M5/114.15.2, Invitrogen, 11-5321-82, 2442242, 1:100 dilution). For Panel 2: CD45.2 BV711 (clone 104, BioLegend, 109847, B348415, 1:100 dilution), CD3 AF488 (clone 17A2, BioLegend, 100210, B364217, 1:100 dilution), CD4 BUV395 (clone GK1.5, BD Biosciences, 563790, 2182147, 1:100 dilution), CD8a PE/Cyanine7 (clone 53-6.7, BioLegend, 100722, B413191, 1:100 dilution), CD44 APC/Cyanine7 (clone IM7, BD Biosciences, 560568, 3128942, 1:100 dilution), CD62L BV421 (clone MEL-14, BD Biosciences, 562910, 3331781, 1:100 dilution), CD69 PerCP/Cyanine5.5 (clone H1.2F3, BioLegend, 104522, B369096, 1:100 dilution), CD279 (PD-1) BV510 (clone 29F.1A12, BioLegend, 135241, B342120, 1:100 dilution), CD223 (LAG-3) BV650 (clone C9B7W, BioLegend, 125227, B364214, 1:100 dilution), CD25 BV605 (clone PC61, BioLegend, 102035, B420344, 1:100 dilution), CD153 PE (clone RM153, Invitrogen, 12-1531-82, 2985938, 1:100 dilution), CD183 BV786 (clone CXCR3-173, BD Biosciences, 741032, 3192579, 1:100 dilution), CD195 (CCR5) BUV737 (clone C34-3448, BD Biosciences, 749670, 3159418, 1:100 dilution). Ghost UV 450 Viability Dye (13-0868-T100, Tonbo Biosciences, D0868061022133) was used as viability dye. Flow cytometry was performed on an LSRFortessa instrument (BD Biosciences).

### Cell viability assay

A total of 5000 cells (from E0771, Eph4, MCF7 and MCF10a cell lines) were seeded in 96-well plates. Serial dilutions of dexamethasone (Dex) from 2560 nM to 40 nM were added to cells after overnight culture. Cells were cultured with DEX at 37°C with 5% CO_2_ for 72 h, then MTT (final concentration is 0.5 mg/mL) was added into each well and further incubated for 4 h. DMSO was added into each emptied well and the quantity of formazan is measured at 590 nm absorbance by using the SpectraMax i3x microplate reader (Molecular Devices).

### Statistical Analyses

All data collected were averaged and expressed as mean ± SEM. Statistical significance was taken as ^∗^p < 0.05, ^∗∗^p < 0.01, and ^∗∗∗^p < 0.001, as determined by Pearson’s correlation, Student’s t test, one-way ANOVA or a two-way repeated-measures ANOVA followed by Bonferroni post hoc tests as appropriate. All n values for each experimental group are described in the appropriate figure legend. Statistical analyses were performed in GraphPad Prism 10.1 (Graphpad, Boston, MA) and MATLAB MATLAB R2024a (The MathWorks, Natick, MA). More specifically, photometry analyses was performed using MATLAB 2024 A and Guided Photometry Analysis in Python (GuPPy)^59^ and RNA Seq analyses were performed using the JupyterHub platform along with FastQC, STAR, FeatureCounts and DeSeq tools. Figures were prepared using BioRender for scientific illustrations/ Schematics, GraphPad Prism v10.1, and Illustrator CC 2023 (Adobe).

## Acknowledgements

The authors acknowledge Dr. Rachel Rubino and all animal care staff (LAR) that provided excellent care to the animals used in these studies. We further thank the following CSHL core facilities: Animal & Tissue Imaging, Flow Cytometry, Sequencing Technologies and Analysis, Microscopy, and Histology cores. We further acknowledge Emma Davidson, Adam Kaufmann, and Dr. Austin Ferro for their insights on loose patch electrophysiology and help in optimizing corticosterone ELISAs, Elena Ghiban for her help with the cDNA library preparation and sequencing for bulk RNA-seq, and Dr. Raditya Utama for his expertise and assistance in the JupyterHub pipeline analysis. We further thank Drs. Linda Van Aelst and Shanu George for early help with electrophysiology. Additionally, we acknowledge all members of the Borniger lab who provided valuable feedback on early versions of this manuscript.

## Author Contributions

Hypothesis generation: A.G., Y.W., J.C.B; Data Acquisition: A.G., Y.W., C.Z., L.B., T-L. W., J.G., C.A., J.C.B.; Data Analysis: A.G., Y.W., C.Z., L.B., C.A.; Drafted manuscript: J.C.B., A.G., Y.W.; Edited manuscript: All authors; Constructed figures: A.G., Y.W., C.Z.; Provided critical reagents/resources: J.C.B., C.A., L.C., Supervised work: J.C.B., C.A., L.C.

---

**Extended Data Fig. 1.**
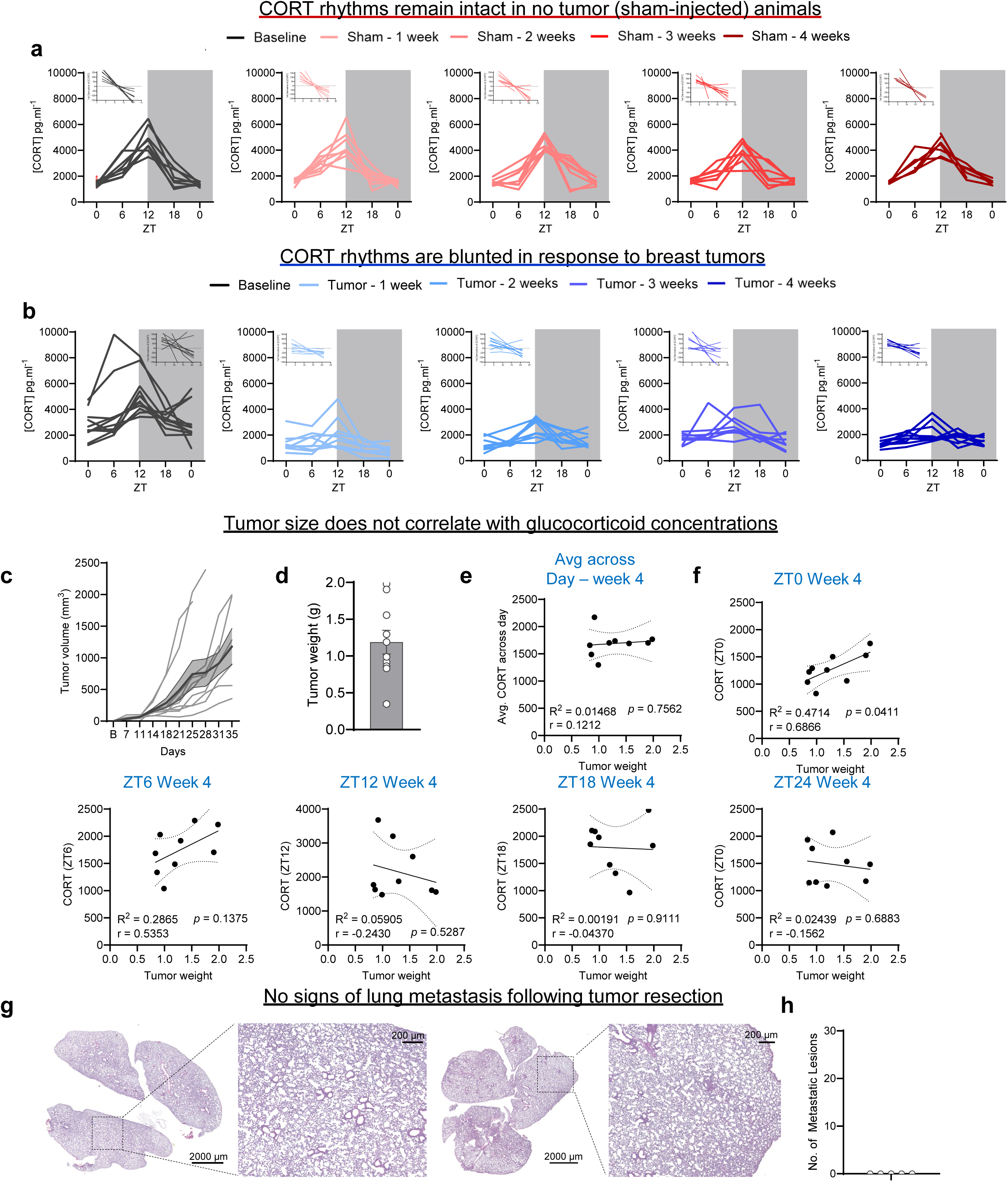
**a,** Individual fecal corticosterone levels across the day (6 hr epochs) in sham-injected mice during baseline and 4 weeks after sham injections. Inset corresponds to the first derivative of individual corticosterone measurements. n = 5-8 mice / timepoint. **b,** Individual fecal corticosterone levels across the day (6 hr epochs) in EO771-injected mice during baseline and 4 weeks of tumor progression. Inset corresponds to the first derivative of individual corticosterone measurements. n = 9-11 mice / timepoint. **c, d,** Tumor progression (**c**) and final tumor weight (**d**) in individual mice across four weeks of tumor progression corresponding to panel **b**. n = 11 mice. **e, f,** Correlation between final tumor weight of EO771-injected mice, and their respective peak corticosterone level at different ZT points measured (**e**) and correlation between final tumor weight and average corticosterone level (average of 5 timepoints) at the 4-week timepoint (**f**). n = 9 mice / correlation. **g,** Representative images showing H&E staining of lungs from mice 9 weeks after breast tumor resection. **h,** Quantification of metastatic lesions in tumor-extracted mice. n = 5 mice. (**j**). Data represented as mean ± SEM.

**Extended Data Fig. 2.**
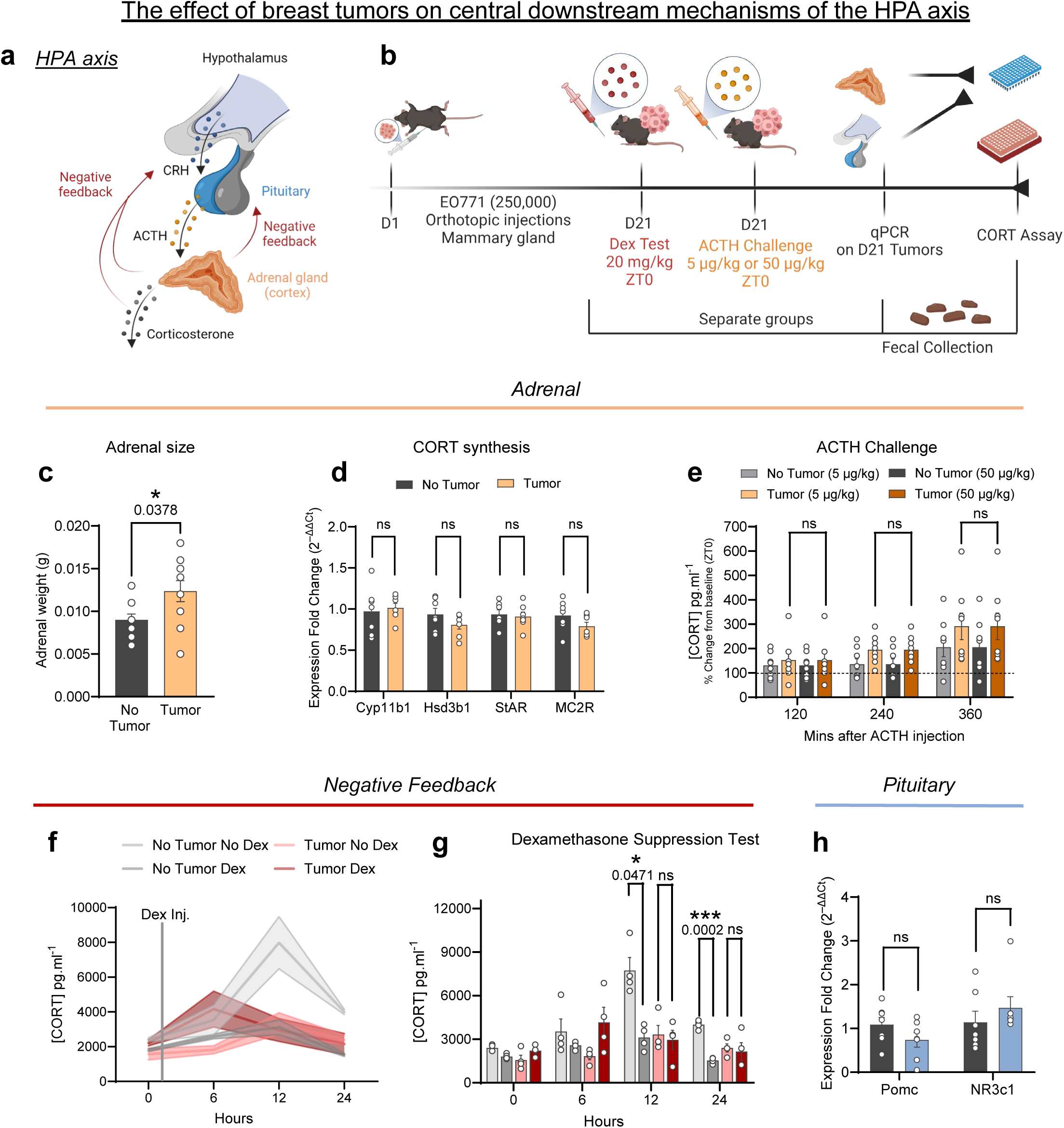
**a,** Schematic highlighting the role in HPA regulation of glucocorticoids. **b,** Schematic depicting corresponding experiments assessing the effect of breast tumors on components of the HPA axis. **c,** Adrenal weights taken from no tumor mice and tumor mice after 4 weeks of tumor growth. n = 9 / no tumor; n = 11 / tumor. **d,** Relative mRNA expression levels of adrenal genes involved with the synthesis glucocorticoids. N = 7 / group. **e,** Fecal corticosterone levels in tumor and non-tumor bearing mice across timepoints (every 2 hrs), after low dose (5 µg/kg, ip, ZT1) and high dose (50 µg/kg, ip, ZT1) injections of an ACTH challenge. n = 8 / group / timepoint. **f, g,** Fecal corticosterone levels across the day in tumor and non-tumor bearing mice during a dexamethasone (10 mg/kg, ip, ZT1) suppression test. n = 4 / group / timepoint. **h,** Relative mRNA expression levels of pituitary genes involved with the synthesis (POMC) of ACTH and glucocorticoid feedback onto the pituitary (NR3c1 / glucocorticoid receptor). n = 7 / group. Data represented as mean ± SEM. \**p* < 0.05, \*\*\**p* < 0.001; two-tailed unpaired student’s t-test (**c, d, h**); two-way repeated-measures ANOVA followed by separate one-way ANOVAs with Bonferroni multiple comparisons (**e-g**).

**Extended Data Fig. 3.**
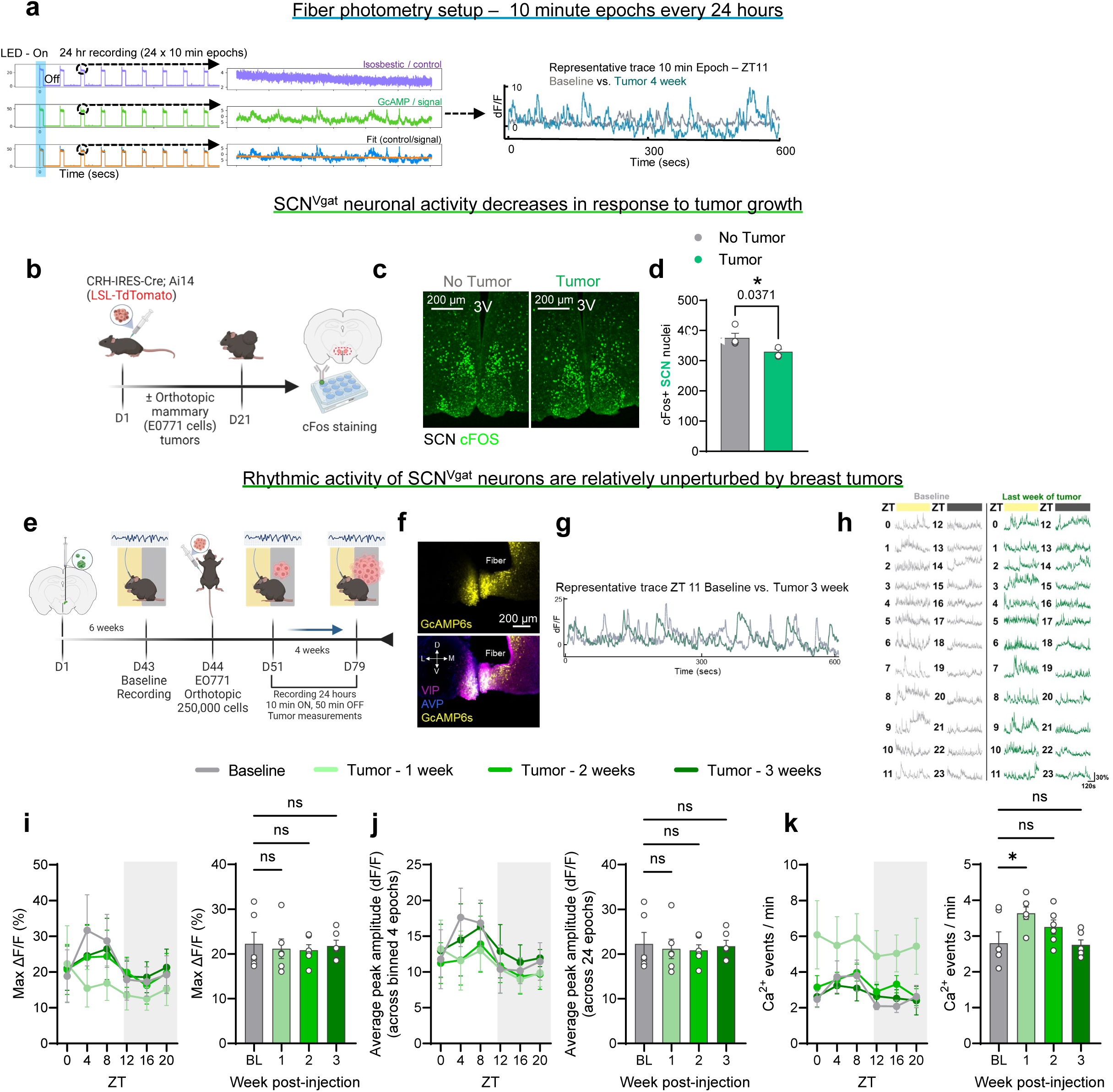
**a,** Schematic of sampling strategy (10 mins / hr for 24 hours) for collecting data regarding fiber photometry recordings during tumor progression. **b,** Timeline for cFos staining in the SCN after sham or EO771 injections. **c,** Representative coronal sections depicting cFos+ (green) nuclei in the SCN in control/ no tumor (left) and tumor (right) mice. **d**, Quantification of cFos+ nuclei in the SCN of no tumor and tumor-bearing mice. n = 4 mice/ group. Each n is average from 2 coronal slices. **e,** Cartoon depicting GCaMP6s viral injection and fiber implantation into the SCN of Vgat-Cre mice and weekly fiber photometry recordings during tumor progression. **f,** Representative coronal section depicting GCaMP6s expression (yellow) within the SCN, and stained with VIP (purple) and AVP (blue) markers. **g-h,** Representative dF/F 10 min epoch trace during a single timepoint (**g**) and during one day (24 hrs) (**h**) of recordings from SCN^Vgat^ neurons between baseline and tumor 3-week timepoints. (**i-k**) Photometry data for weekly recordings (10 min epochs) during tumor progression depicting max dF/F (**i**), average peak amplitude (**j**), and number of calcium transients (**k**) for SCN^Vgat^ neurons collapsed into 4 hr averages (left panels) and total timepoints (right panels), respectively. n = 5 mice per baseline and tumor week. Data represented as mean ± SEM. \**p* < 0.05; two-tailed unpaired student’s t-test (**d**); two-way repeated-measures ANOVA followed by separate one-way ANOVAs with Bonferroni multiple comparisons (**i-k**).

**Extended Data Fig. 4.**
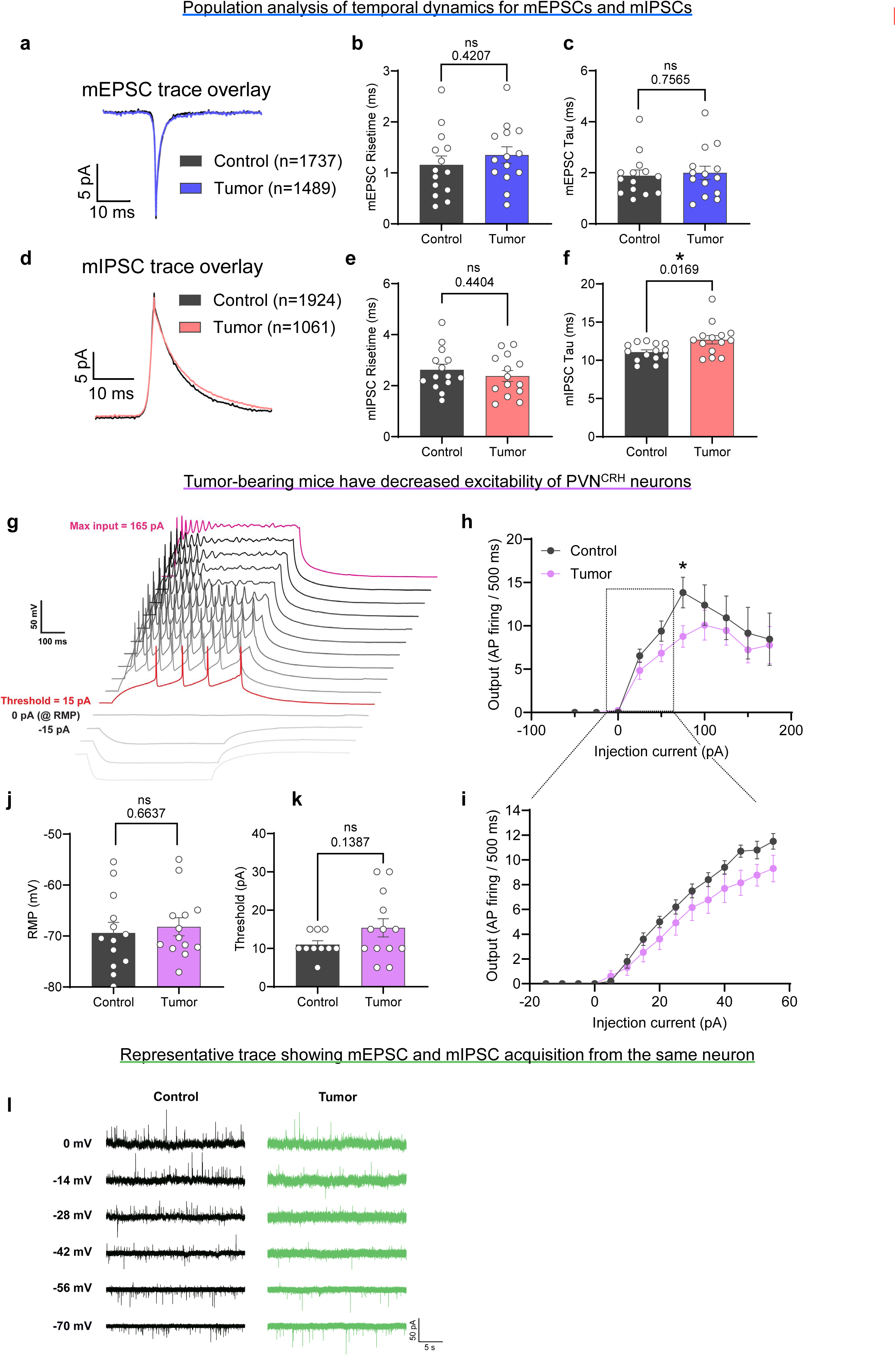
**a,** Overlay of the mean mEPSC traces of PVN^CRH^ cells from control (black) and tumor (blue) mice. **b-c,** Population comparison of the mEPSC temporal dynamics showing no significant changes observed for risetime (**b**) or decay (**c**). **d-f** Overlay of the mean mIPSC traces showing no significant difference in inhibition activation dynamics **(e),** but a longer decay was observed in PVN^CRH^ neurons from tumor-bearing mice (**f**). **g,** Example traces from current-clamp experiment showing voltage changes of one PVN^CRH^ neuron in response to injecting currents in +15 pA steps for 500 ms. The resting membrane potential was recorded at 0 pA step, and the minimal current level that elicits action potentials is considered the threshold of the neuron (red trace). **h,** Population action potential numbers are processed offline and plotted as the input-output function. n=13 from 4 mice for control and n=13 from 3 mice for tumor. The PVN^CRH^ neurons in tumor-bearing mice exhibit decreasing excitability as the input increases within the dynamic range, with the difference reaches significant level near neurons’ maximum firing capability. **i,** A current clamp protocol with +5 pA steps were added at para-threshold levels for a more accurate acquisition of the threshold. n=10 from 3 mice for control and n=13 from 3 mice for tumor. **j, k,** Together with the observation that no significant difference was observed in resting membrane potential (**j**) or threshold (**k**), this result suggests a potential influence on the gain-controlling mechanism of the PVN^CRH^ neurons by breast tumor. n=13 from 4 mice for control and n=13 from 3 mice for tumor in **j**, and n=10 from 3 mice for control and n=13 from 3 mice for tumor in **k**. **l,** Representative traces of the E/I experiment from control (left) and tumor-bearing (right) mouse. Note the gradual deactivation of the downward, excitatory currents and the activation of the upward, inhibitory currents as the neurons were held at increasingly depolarizing voltages from −70 mV to 0 mV. Data represented as mean ± SEM. \**p* < 0.05; two-tailed unpaired student’s t-test (**b, c, e, f, j, k**); two-way repeated-measures ANOVA followed by separate one-way ANOVAs with Bonferroni multiple comparisons (**h, i**).

**Extended Data Fig. 5.**
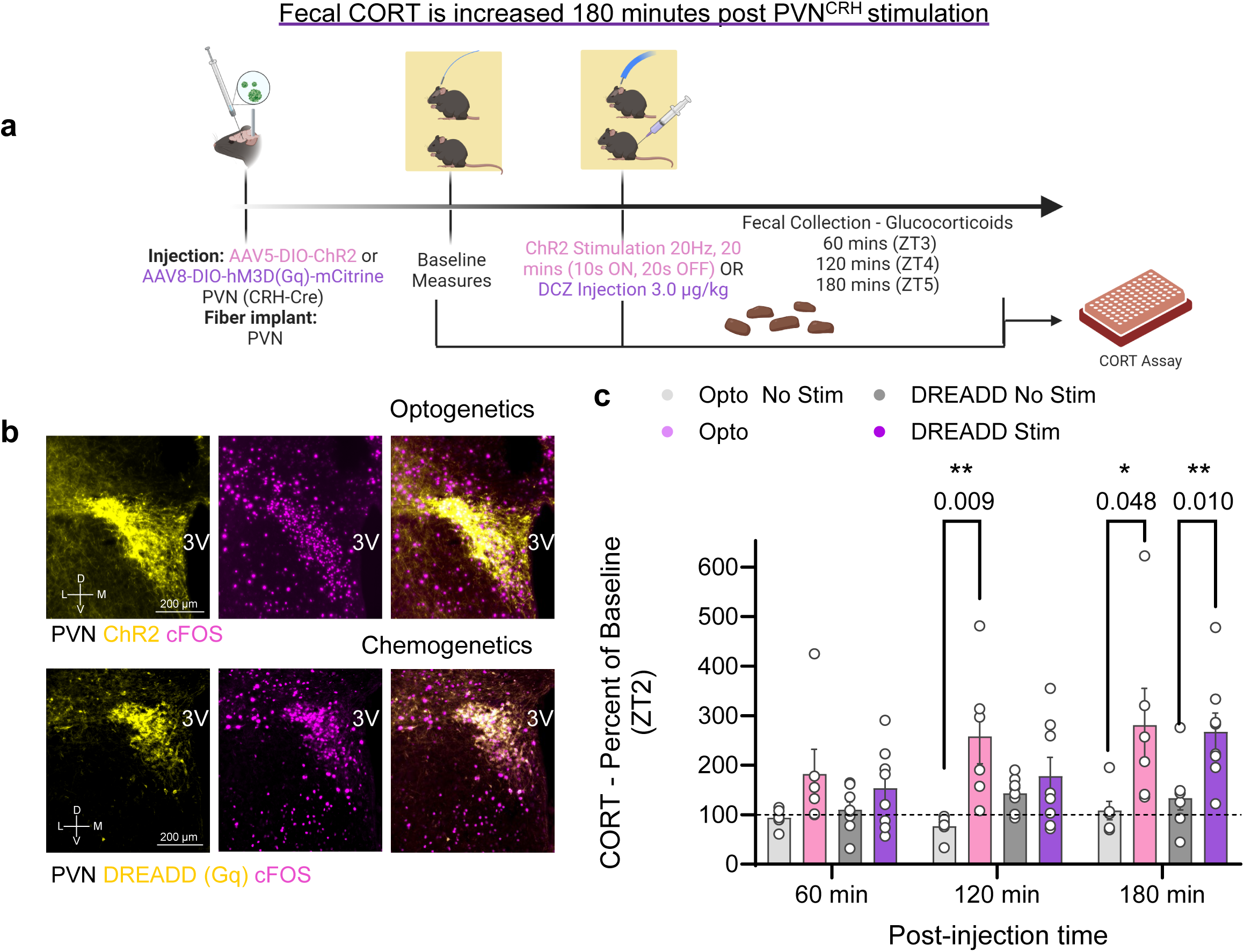
**a,** Schematic depicting fecal collection following ChR2-mediated stimulation (optogenetics) or DCZ-mediated stimulation (DREADDs) of PVN^CRH^ neurons, to determine when increases in corticosterone are observed in fecal boli post stimulation. **b,** Coronal sections depicting ChR2 (yellow, top), DREADDs (yellow, bottom) and cFOS (purple) expression within the PVN of CRH-Cre+ mice after either blue light (473 nm) or DCZ (3 µg/kg, ip) stimulation. **c,** Hourly fecal corticosterone measurements expressed as a percent of baseline (ZT2) after no stimulation, 20 Hz opto-stimulation, and DREADD-mediated stimulation (DCZ, 3 µg/kg, ip) of PVN^CRH^ neurons. n = 6 / opto groups; n = 8 / DREADD groups Data represented as mean ± SEM. \**p* < 0.05, \*\**p* < 0.01; two-way repeated-measures ANOVA followed by separate one-way ANOVAs with Bonferroni multiple comparisons (**c**).

**Extended Data Fig. 6.**
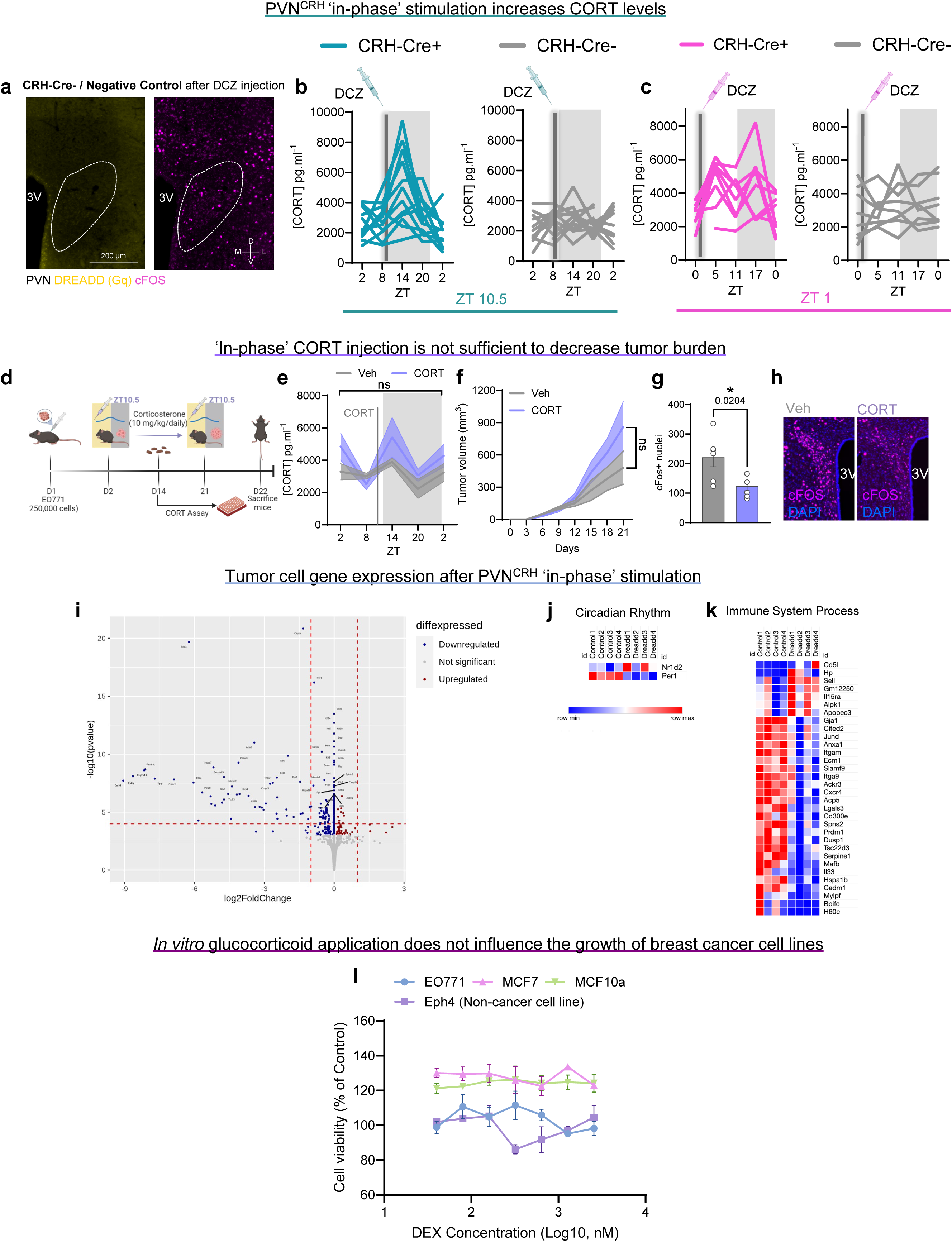
**a,** Representative coronal section depicting DREADD (Gq, yellow) and cFOS (purple) expression within the PVN of CRH-Cre-(control) mouse after DCZ injection. **b, c,** Individual measurements of corticosterone levels in CRH-Cre+ and CRH-Cre-mice across the day in ZT10.5 injected mice (**b**) and ZT1 injected mice (**c**). n = 8 Cre- and 7-8 Cre+ mice / timepoint for ZT1; n = 10-12 CRH-Cre- and 11-12 CRH-Cre+ mice / timepoint for ZT10.5. **d,** Schematic depicting daily, corticosterone (10 mg/kg, ip) injections at ZT10.5 of mice during tumor progression. **e,** Individual fecal corticosterone measurements across day 14 of tumor progression during a ZT10.5 CORT or vehicle injection. n = 5 mice / group. **f,** Progression of mammary tumor size during 21 days after initial tumor injection from the ZT10.5 CORT-injected mice. n = 5 mice / group. **g,** Quantification of cFOS+ nuclei in the PVN of mice after the last ZT10.5 CORT or vehicle injection. n = 5 mice / group. Each mouse represents an average from 2 coronal slices. **h,** Coronal sections depicting cFOS (purple) expression and DAPI (blue) within the PVN of mice after the last ZT10.5 CORT or vehicle injection. **i**, Volcano plot showing differential gene expression analysis from the bulk RNA-seq of “in-phase” stimulated and control groups. Each point represents a single gene, with the x-axis showing the log2 fold change in expression between “in-phase” stimulated group vs. controls, and the y-axis displaying the -log10 adjusted p-value. Heatmap of genes enriched in Circadian Rhythm (**j**) and Immune System Process (**k**) GO term categories with 1.2-fold change cutoff. n = 4 mice / group. **l,** MTT assay assessing cell viability in breast cancer cell lines EO771 (mouse), MCF7 (human), and epithelial cell lines Eph4 (mouse), MCF10a (human) treated with serial concentrations of dexamethasone. n = 3 replicates / cell line. All data represented as mean ± SEM. \**p* < 0.05. Two-tailed unpaired student’s t-test (**g**); two-way repeated-measures ANOVA followed by separate one-way ANOVAs with Bonferroni multiple comparisons (**e, f, l**).

**Extended Data Fig. 7.**
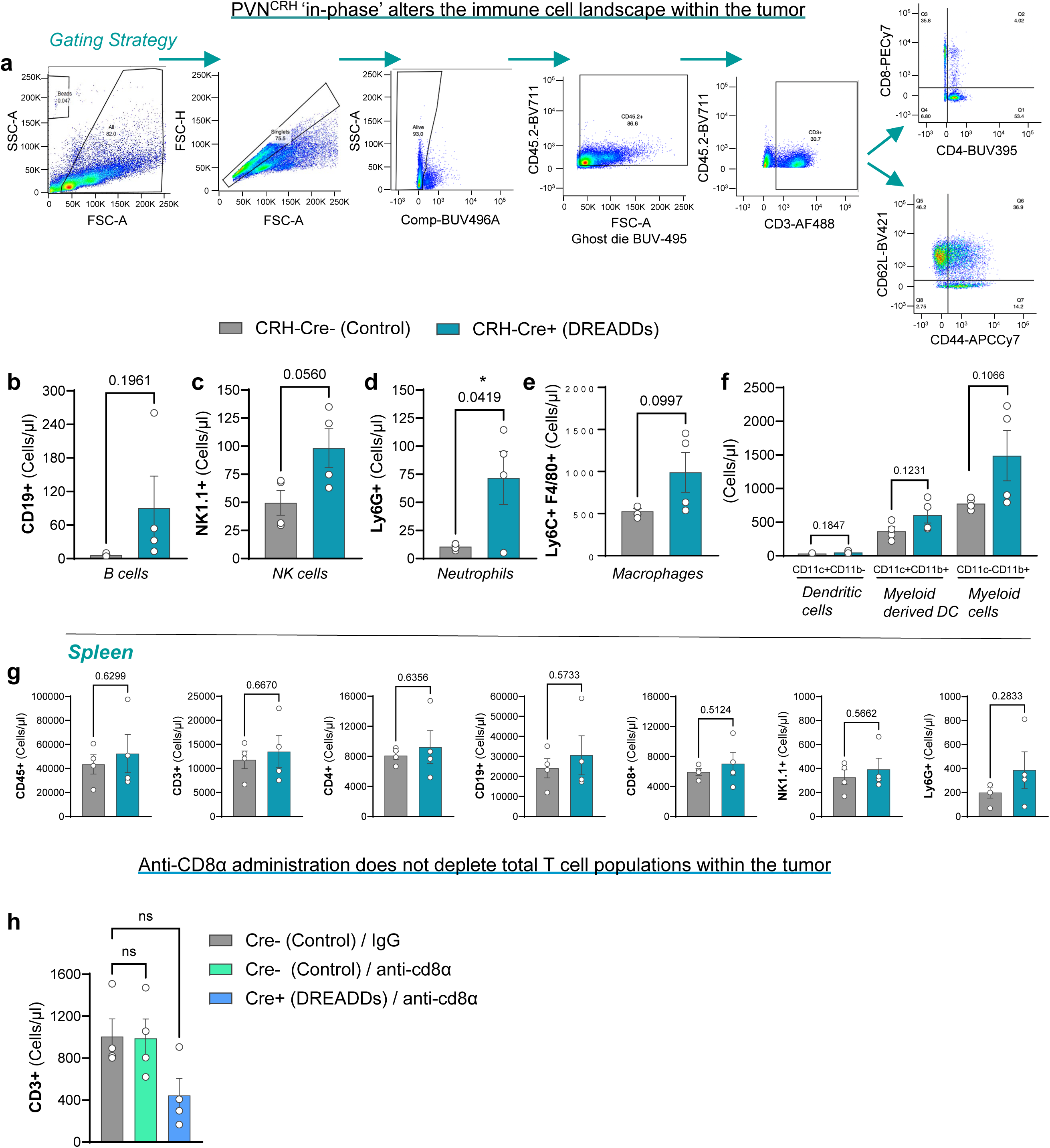
**a,** Gating strategy for flow cytometric analysis of tumor immune cell markers in the ZT10.5-treated DCZ groups. **b-f,** Quantification of CD19+ (**b**), NK1.1+ (**c**), Ly6G+ (**d**), Ly6C+ F4/80+ (**e**), and CD11cCD11b (**f**) cells of CRH-Cre- and CRH-Cre+ breast tumors after daily CRH stimulation at ZT10.5 and analyzed by flow cytometry. n = 4 mice / group. **g,** Quantification of CD45+ (**b**), CD3+, CD4+, CD19+, CD8+, NK1.1+, and Ly6G+ from spleen in ZT10.5 DCZ-treated mice. n = 4 mice / group. **h,** Quantification of CD3+ breast tumor cells in ZT10.5 DCZ-injected mice after IgG control or CD8+ depletion and analyzed by flow cytometry. n = 4 mice / group. All data represented as mean ± SEM. \**p* < 0.05; two-tailed unpaired student’s t-test (**b-g**); one-way ANOVAs with Bonferroni multiple comparisons (**h**).

## References

1. Sephton, S. E., Sapolsky, R. M., Kraemer, H. C. & Spiegel, D. Diurnal Cortisol Rhythm as a Predictor of Breast Cancer Survival. JNCI J. Natl. Cancer Inst. 92, 994–1000 (2000).

2. Abercrombie, H. C. et al. Flattened cortisol rhythms in metastatic breast cancer patients. Psychoneuroendocrinology 29, 1082–1092 (2004).

3. Bower, J. E. et al. Diurnal cortisol rhythm and fatigue in breast cancer survivors. Psychoneuroendocrinology 30, 92–100 (2005).

4. DeSantis, C. E. et al. International Variation in Female Breast Cancer Incidence and Mortality Rates. Cancer Epidemiol. Biomarkers Prev. 24, 1495–1506 (2015).

5. Ancoli-Israel, S. et al. Fatigue, sleep, and circadian rhythms prior to chemotherapy for breast cancer. Support. Care Cancer 14, 201–209 (2006).

6. Ancoli-Israel, S., Moore, P. J. & Jones, V. The relationship between fatigue and sleep in cancer patients: A review. Eur. J. Cancer Care (Engl.) 10, 245–255 (2001).

7. Berger, A. M., Farr, L. A., Kuhn, B. R., Fischer, P. & Agrawal, S. Values of Sleep/Wake, Activity/Rest, Circadian Rhythms, and Fatigue Prior to Adjuvant Breast Cancer Chemotherapy. J. Pain Symptom Manage. 33, 398–409 (2007).

8. Schmidt, M. E. et al. Cancer-related fatigue shows a stable association with diurnal cortisol dysregulation in breast cancer patients. Brain. Behav. Immun. 52, 98–105 (2016).

9. Sephton, S. E. et al. Depression, cortisol, and suppressed cell-mediated immunity in metastatic breast cancer. Brain. Behav. Immun. 23, 1148–1155 (2009).

10. Hsiao, F.-H. et al. A longitudinal study of diurnal cortisol patterns and associated factors in breast cancer patients from the transition stage of the end of active cancer treatment to post-treatment survivorship. Breast Edinb. Scotl. 36, 96–101 (2017).

11. Sephton, S. E. et al. Diurnal cortisol rhythm as a predictor of lung cancer survival. Brain. Behav. Immun. 30 Suppl, S163-170 (2013).

12. Rich, T. et al. Elevated Serum Cytokines Correlated with Altered Behavior, Serum Cortisol Rhythm, and Dampened 24-Hour Rest-Activity Patterns in Patients with Metastatic Colorectal Cancer. Clin. Cancer Res. 11, 1757–1764 (2005).

13. Schrepf, A. et al. Diurnal cortisol and survival in epithelial ovarian cancer. Psychoneuroendocrinology 53, 256–267 (2015).

14. Shimba, A. et al. Glucocorticoids Drive Diurnal Oscillations in T Cell Distribution and Responses by Inducing Interleukin-7 Receptor and CXCR4. Immunity 48, 286–298.e6 (2018).

15. So, A. Y.-L., Bernal, T. U., Pillsbury, M. L., Yamamoto, K. R. & Feldman, B. J. Glucocorticoid regulation of the circadian clock modulates glucose homeostasis. Proc. Natl. Acad. Sci. 106, 17582–17587 (2009).

16. Balsalobre, A. et al. Resetting of Circadian Time in Peripheral Tissues by Glucocorticoid Signaling. Science 289, 2344–2347 (2000).

17. Oster, H. et al. The Functional and Clinical Significance of the 24-Hour Rhythm of Circulating Glucocorticoids. Endocr. Rev. 38, 3–45 (2017).

18. Jones, J. R., Chaturvedi, S., Granados-Fuentes, D. & Herzog, E. D. Circadian neurons in the paraventricular nucleus entrain and sustain daily rhythms in glucocorticoids. Nat. Commun. 12, 5763 (2021).

19. Oster, H. et al. The circadian rhythm of glucocorticoids is regulated by a gating mechanism residing in the adrenal cortical clock. Cell Metab. 4, 163–173 (2006).

20. Le Naour, A. et al. EO771, the first luminal B mammary cancer cell line from C57BL/6 mice. Cancer Cell Int. 20, 328 (2020).

21. Touma, C., Palme, R. & Sachser, N. Analyzing corticosterone metabolites in fecal samples of mice: a noninvasive technique to monitor stress hormones. Horm. Behav. 45, 10–22 (2004).

22. Boyle, S. T., Faulkner, J. W., McColl, S. R. & Kochetkova, M. The chemokine receptor CCR6 facilitates the onset of mammary neoplasia in the MMTV-PyMT mouse model via recruitment of tumor-promoting macrophages. Mol. Cancer 14, 115 (2015).

23. Gjerstad, J. K., Lightman, S. L. & Spiga, F. Role of glucocorticoid negative feedback in the regulation of HPA axis pulsatility. Stress 21, 403–416 (2018).

24. Spencer, R. L. & Deak, T. A USERS GUIDE TO HPA AXIS RESEARCH. Physiol. Behav. 178, 43–65 (2017).

25. Hill, M. N. & Tasker, J. G. Endocannabinoid signaling, glucocorticoid-mediated negative feedback, and regulation of the hypothalamic-pituitary-adrenal axis. Neuroscience 204, 5–16 (2012).

26. Nagai, Y. et al. Deschloroclozapine, a potent and selective chemogenetic actuator enables rapid neuronal and behavioral modulations in mice and monkeys. Nat. Neurosci. 23, 1157–1167 (2020).

27. Li, S.-B. et al. Hypothalamic circuitry underlying stress-induced insomnia and peripheral immunosuppression. Sci. Adv. 6, eabc2590 (2020).

28. Poller, W. C. et al. Brain motor and fear circuits regulate leukocytes during acute stress. Nature 607, 578– 584 (2022).

29. Guo, Y., Luan, L., Patil, N. K. & Sherwood, E. R. Immunobiology of the IL-15-IL-15Rα Complex as an Antitumor and Antiviral Agent. Cytokine Growth Factor Rev. 38, 10–21 (2017).

30. Huntoon, K. M. et al. The acute phase protein haptoglobin regulates host immunity. J. Leukoc. Biol. 84, 170–181 (2008).

31. Sanjurjo, L., Aran, G., Roher, N., Valledor, A. F. & Sarrias, M.-R. AIM/CD5L: a key protein in the control of immune homeostasis and inflammatory disease. J. Leukoc. Biol. 98, 173–184 (2015).

32. Fortunato, O. et al. CXCR4 Inhibition Counteracts Immunosuppressive Properties of Metastatic NSCLC Stem Cells. Front. Immunol. 11, 02168 (2020).

33. Chen, F. et al. Histone deacetylase 2 regulates STAT1-dependent upregulation of atypical chemokine receptor 3 to induce M2 macrophage migration and immune escape in hepatocellular carcinoma. Mol. Immunol. 151, 204–217 (2022).

34. Choi, E. Y., Choi, K., Nam, G., Kim, W. & Chung, M. H60: A Unique Murine Hematopoietic Cell-Restricted Minor Histocompatibility Antigen for Graft-versus-Leukemia Effect. Front. Immunol. 11, (2020).

35. Ameri, A. H. et al. IL-33/regulatory T cell axis triggers the development of a tumor-promoting immune environment in chronic inflammation. Proc. Natl. Acad. Sci. U. S. A. 116, 2646–2651 (2019).

36. Vega, M. R. de la. A holistic view of cancer. Cancer Cell 41, 373 (2023).

37. Hirata, T. Axon Outgrowth. in Encyclopedia of Neuroscience (eds. Binder, M. D., Hirokawa, N. & Windhorst, U.) 311–313 (Springer, Berlin, Heidelberg, 2009). doi:10.1007/978-3-540-29678-2_515.

38. Gerendai, I. et al. Transneuronal labelling of nerve cells in the CNS of female rat from the mammary gland by viral tracing technique. Neuroscience 108, 103–118 (2001).

39. Köves, K., Györgyi, Z., Szabó, F. K. & Boldogkői, Z. Characterization of the autonomic innervation of mammary gland in lactating rats studied by retrograde transynaptic virus labeling and immunohistochemistry. Acta Physiol. Hung. 99, 148–158 (2012).

40. Perkinson, M. R., Kim, J. S., Iremonger, K. J. & Brown, C. H. Visualising oxytocin neurone activity in vivo: The key to unlocking central regulation of parturition and lactation. J. Neuroendocrinol. 33, e13012 (2021).

41. Pan, S. et al. Stimulation of hypothalamic oxytocin neurons suppresses colorectal cancer progression in mice. eLife 10, e67535 (2021).

42. Vallières, L. & Rivest, S. Interleukin-6 Is a Needed Proinflammatory Cytokine in the Prolonged Neural Activity and Transcriptional Activation of Corticotropin-Releasing Factor during Endotoxemia1. Endocrinology 140, 3890–3903 (1999).

43. Sapolsky, R., Rivier, C., Yamamoto, G., Plotsky, P. & Vale, W. Interleukin-1 stimulates the secretion of hypothalamic corticotropin-releasing factor. Science 238, 522–524 (1987).

44. Berkenbosch, F., Oers, J. van, Rey, A. del, Tilders, F. & Besedovsky, H. Corticotropin-releasing factor-producing neurons in the rat activated by interleukin-1. Science 238, 524–526 (1987).

45. Schmidt, E. D., Janszen, A. W., Wouterlood, F. G. & Tilders, F. J. Interleukin-1-induced long-lasting changes in hypothalamic corticotropin-releasing hormone (CRH)--neurons and hyperresponsiveness of the hypothalamus-pituitary-adrenal axis. J. Neurosci. 15, 7417–7426 (1995).

46. Khazaeipool, Z., Wiederman, M. & Inoue, W. Prostaglandin E2 depresses GABA release onto parvocellular neuroendocrine neurones in the paraventricular nucleus of the hypothalamus via presynaptic receptors. J. Neuroendocrinol. 30, e12638 (2018).

47. Sun, Q. et al. Area postrema neurons mediate interleukin-6 function in cancer cachexia. Nat. Commun. 15, 4682 (2024).

48. Boudaba, C., Szabó, K. & Tasker, J. G. Physiological Mapping of Local Inhibitory Inputs to the Hypothalamic Paraventricular Nucleus. J. Neurosci. 16, 7151–7160 (1996).

49. Chou, T. C. et al. Critical Role of Dorsomedial Hypothalamic Nucleus in a Wide Range of Behavioral Circadian Rhythms. J. Neurosci. 23, 10691–10702 (2003).

50. Yoshida, S. et al. Elucidation of the mechanisms underlying tumor aggravation by the activation of stress-related neurons in the paraventricular nucleus of the hypothalamus. Mol. Brain 16, 18 (2023).

51. Nelson, R. J., DeVries, A. C. & Prendergast, B. J. Researchers need to better addre ss time-of-day as a critical biological variable. Proc. Natl. Acad. Sci. 121, e2316959121 (2024).

52. Pick, R., He, W., Chen, C.-S. & Scheiermann, C. Time-of-Day-Dependent Trafficking and Function of Leukocyte Subsets. Trends Immunol. 40, 524–537 (2019).

53. Pick, R., Wang, C., Zeng, Q., Gül, Z. M. & Scheiermann, C. Circadian Rhythms in Anticancer Immunity: Mechanisms and Treatment Opportunities. Annu. Rev. Immunol. 42, 83–102 (2024).

54. Obradović, M. M. S. et al. Glucocorticoids promote breast cancer metastasis. Nature 1 (2019) doi:10.1038/s41586-019-1019-4.

55. Rasiah, N. P., Loewen, S. P. & Bains, J. S. Windows into stress: a glimpse at emerging roles for CRHPVN neurons. Physiol. Rev. 103, 1667–1691 (2023).

56. Jansen, A. S. P., Van Nguyen, X., Karpitskiy, V., Mettenleiter, T. C. & Loewy, A. D. Central Command Neurons of the Sympathetic Nervous System: Basis of the Fight-or-Flight Response. Science 270, 644– 646 (1995).

57. Hosoi, T., Okuma, Y. & Nomura, Y. Electrical stimulation of afferent vagus nerve induces IL-1β expression in the brain and activates HPA axis. Am. J. Physiol.-Regul. Integr. Comp. Physiol. 279, R141–R147 (2000).

58. Clancy, J. A. et al. Non-invasive Vagus Nerve Stimulation in Healthy Humans Reduces Sympathetic Nerve Activity. Brain Stimulat. 7, 871–877 (2014).

59. Sherathiya, V. N., Schaid, M. D., Seiler, J. L., Lopez, G. C. & Lerner, T. N. GuPPy, a Python toolbox for the analysis of fiber photometry data. Sci. Rep. 11, 24212 (2021).

60. Tomayko, M. M. & Reynolds, C. P. Determination of subcutaneous tumor size in athymic (nude) mice. Cancer Chemother. Pharmacol. 24, 148–154 (1989).

61. Konno, K., Yamasaki, M., Miyazaki, T. & Watanabe, M. Glyoxal fixation: An approach to solve immunohistochemical problem in neuroscience research. Sci. Adv. 9, eadf7084 (2023).

